# Unmasking Pathogen Traits for Chronic Colonization in Neurogenic Bladder Patients

**DOI:** 10.1101/2025.08.14.669717

**Authors:** Seth A. Reasoner, Brendan T. Frainey, Owen F. Hale, Alexandra Borden, M. Kyle Graham, Elise Turner, Lucas R. Brenes, Carl B.W. Soderstrom, Hamilton Green, Jonathan E. Schmitz, Michael T. Laub, Maryellen S. Kelly, Douglass B. Clayton, Maria Hadjifrangiskou

**Author notes:** These authors contributed equally to this work* (Seth A. Reasoner, Brendan T. Frainey). **CORRESPONDING AUTHORS:** Brendan T. Frainey, MD, Maria Hadjifrangiskou, PhD.

## Abstract

Individuals with neurogenic bladder are particularly susceptible to both chronic bacterial colonization of the bladder and urinary tract infections (UTIs). Neurogenic bladder can arise from a variety of diseases such as diabetes, spinal cord injuries, and spina bifida. To study the ecological and evolutionary dynamics of the microbiome in neurogenic bladder, we developed a longitudinal cohort of 77 children and young adults with spina bifida from two medical centers. We used enhanced urine culture, 16S rRNA sequencing, and whole genome sequencing to characterize the microbial composition of urine and fecal samples. In addition to prospective sample collection, we retrieved prior bacterial isolates from enrolled patients from Vanderbilt’s clinical microbial biobank, MicroVU. This allowed us to compare bacterial isolates from the same patients over a period of five years. Urine samples were characterized by high abundance of urinary pathogens, such as *E. coli* and *Klebsiella*. From longitudinal isolates from individual patients, we identified two common patterns of urinary tract colonization. We observed either the rapid cycling of strains and/or species, often following antibiotic treatment, or we observed the persistence of a single strain across timepoints. Neither persistence of a strain nor colonization with a new strain or species was associated with increased antibiotic resistance. Rather, in paired longitudinally collected strains from the same patients, mutations were identified in genes that code for cell envelope components associated with immune or phage evasion. Experimental testing revealed that O-antigen/LPS biosynthesis mutations confer protection from the immune system while altering susceptibility to phage predation, reflecting a fitness trade-off. We argue that this unparalleled cohort offers the opportunity to identify mechanisms of bacterial adaptation to the urinary tract that can be exploited in future therapeutic approaches.

## 1. INTRODUCTION

Individuals with neurogenic bladder are particularly susceptible to chronic bacterial colonization of the bladder and urinary tract infections (UTIs)^1^. Neurogenic bladder encompasses a broad range of neurologic conditions that impair innervation and function of the bladder. Examples of diseases that can lead to mild or moderate neurogenic bladder include multiple sclerosis and neuropathy from diabetes^2,3^. Two of the most severe forms of neurogenic bladder are the result of spina bifida and spinal cord injuries. The severity of spina bifida and spinal cord injuries leads to multiple risk factors for urinary tract bacterial colonization; these risk factors include urinary stasis due to incomplete/poor bladder emptying, routine urinary tract instrumentation, and concomitant neurogenic bowel. Improper bladder contraction often requires that these afflicted individuals self-catheterize to relieve bladder pressure.

We chose to use spina bifida as a model disease to understand the ecological dynamics of the urinary microbiome and UTIs in neurogenic bladder. Spina bifida is a congenital malformation of the spinal cord, arising from incomplete embryonic closure of the neural tube. Spina bifida affects multiple organ systems, including neurogenic bladder and bowel, hydrocephalus, orthopedic abnormalities of the spinal cord, and reduced lower extremity function^4,5^. Urologic issues, such as urinary incontinence, urolithiasis, and recurrent UTIs, have a large impact on the lives of individuals with spina bifida^6^. Specifically, UTIs are a common reason for hospitalization, morbidity, and mortality in these patients^7–10^.

The diagnosis of UTI in patients with spina bifida, and neurogenic bladder broadly, is complicated by reduced or absent pelvic sensation intrinsic to neurogenic bladder pathophysiology^11,12^. The diagnosis of UTI requires patient symptoms in addition to evidence of bacteria in the urine^13^. Patients with spina bifida are frequently colonized with pathogenic bacteria yet unable to feel irritative symptoms of UTI. The inability to feel local symptoms of UTI often leads to bacteria spreading to the upper urinary tract (i.e., kidneys) resulting in pyelonephritis and sometimes. Thus, clinicians are apt to treat these patients aggressively with antibiotics to prevent severe infections. Understanding and improving the differentiation of urinary tract bacterial colonization from infection is a vital endeavor to improve the care of patients with spina bifida and neurogenic bladder.

In this study, we sought to characterize the urinary and intestinal microbiomes over time in children with spina bifida. In addition to sequencing microbial communities, we performed whole-genome sequencing on individual bacterial isolates from urine samples. We utilized our medical center’s retrospective microbial biobank MicroVU to compare previously collected urinary bacterial isolates to contemporary isolates from the same patient to understand what genomic changes accumulate over time that may facilitate pathogen persistence. We hypothesized that selective pressures would drive the evolution of pathogen-associated genes, enabling more effective bacterial colonization and persistence in the urinary tract. From longitudinally collected samples, we identified mutations in genes encoding bacterial envelope-associated components, including siderophore receptors, two-component signal transduction systems, as well as in LPS and O-antigen biosynthesis genes. We detected no changes in antibiotic resistance carriage in this study. Rather, we observe enrichment of mutations that indicate remodeling of the pathogen cell envelope towards a less immunogenic state. We provide *in vitro* experimental evidence that these mutations promote immune evasion and do not impair bacterial fitness in a murine model of bladder infection. However, we provide evidence that this cell envelope remodeling leads to altered susceptibility of the pathogen to phage; an avenue that could be explored as a potential personalized therapeutic approach. This dataset represents a unique longitudinal resource for identifying bacterial pathogen evolution in the urinary tract and advancing mechanistic research about bacterial adaptation to the urinary tract.

## 2. METHODS

### 2.1 Patient Enrollment, Sample and Metadata Collection

This study was approved by the Vanderbilt University Medical Center Institutional Review Board (IRB # 191815) and Duke University Health System Institutional Review Board (IRB # 00114763). Subjects ages 0 to 30 years of age were included in the study. The parents or guardians of subjects under 18 provided consent to participate in the study, and children ages 7 to 17 years old provided assent. Subjects were enrolled in either a multidisciplinary outpatient spina bifida clinic at the time of urodynamic testing or after birth within the neonatal intensive care unit (NICU). Cohort data is provided in **Supplementary Table 1**.

Urine and fecal samples were collected during routine urodynamic testing. Catheterized urine was obtained using a sterile technique either per urethra or via a catheterizable channel if present. Fecal material was either obtained from formed stool present in the diaper at the time of urodynamics or from the urodynamic rectal catheter tip which was vortexed in PBS. Hand swabs were collected from the primary individual responsible for catheterizing (patient or caregiver) if the patient was on clean intermittent catheterization. Hands were swabbed with a sterile swab (Fisher, 1490710) for 1 minute. In newborn patients admitted to the NICU, catheterized urine specimens were obtained on day of life (DOL) 0 prior to antibiotics. Meconium samples were obtained from newborns after their first stool. All samples were stored at -80 °C until DNA extraction.

Retrospective bacterial isolates were retrieved from the microbial biobank MicroVU. MicroVU stores a random subset of all clinical bacterial isolates from the Vanderbilt University Medical Center Clinical Microbiology laboratory. Bacterial isolates were identified by unique patient identifiers by the VUMC Integrated Data Access and Services Core (IDASC). Isolates were matched to documented urine cultures in the patient’s medical record.

### 2.2 Sample Nomenclature

The first string of numbers is the patient ID. The type of sample is denoted by the first number after the dash (-), 1: urine; 2: fecal; 3: hand swab. The timepoint at which the sample was collected is denoted by the second number after the dash (e.g., BTF1-23, subject 1, fecal specimen, timepoint 3) (**Supplementary Table 2**). Whole genome sequenced isolates only came from urine samples (**Supplementary Table 3**). They are denoted by the patient ID, followed by a “T” for timepoint. Prospectively collected samples have positive integers following the “T” (e.g., BTF1T1), whereas retrospective isolates from the MicroVU biobank have negative integers following the “T” (e.g., BTF1T-1).

### 2.3 Enhanced Urine Culture

For samples collected at Vanderbilt, urine samples were plated within 4 hours of collection. Urine samples from Duke University were placed in boric acid preservative tubes (BD 364951) and shipped overnight to Vanderbilt. Urine samples were plated on Brucella, blood, and MacConkey agar plates in duplicate, 100 µL on each plate. One set of plates was incubated in a 5% CO_2_ atmosphere at 37°C and another set of plates was incubated at 37°C anaerobically (BD GasPak™ EZ anaerobe pouch). The GasPak™ EZ anaerobe pouch contains a methylene blue indicator tablet which is colorless under anaerobic conditions. Following 48-72 hours of growth, the biomass was scraped off the plate and stored in BHI-glycerol at -80 °C.

### 2.4 Antimicrobial Susceptibility Testing

Antimicrobial susceptibility was tested by broth microdilution using 2-fold dilutions of the following antibiotics: ampicillin (0-256 µg/mL), ceftriaxone (0-64 µg/mL), ciprofloxacin (0-8 µg/mL), gentamicin (0-64 µg/mL), nitrofurantoin (0-256 µg/mL), tetracycline (0-64 µg/mL), and trimethoprim-sulfamethoxazole (0-32/608 µg/mL). Minimum inhibitory concentrations (MIC) are reported in **Supplementary Table 4**.

### 2.5 16S rRNA Sequencing and Bioinformatic Analysis of 16S rRNA Sequencing Data

The first batch of samples was processed at University of California San Diego Microbiome Core, and the second batch of samples was processed at Argonne National Lab. DNA was extracted from 0.5mL of urine, fecal suspensions, and swabs using the MagMAX™ Microbiome Ultra Nucleic Acid Isolation Kit (A42357, ThermoFisher) in batch one, and the DNeasy PowerSoil kit (47014, Qiagen) in batch two. Ten-fold dilutions of the ZymoBIOMICS™ Microbial Community Standard (D6300, Zymo) were extracted alongside the patient samples and included in both batches to benchmark differences in batches. Extraction blanks were included on each plate of DNA extraction. The V4 region of the 16S rRNA gene was amplified with the 515F and 806R primers from the Earth Microbiome Project^14^. Paired-end 150 bp reads were sequenced by Illumina MiSeq chemistry.

Raw 16S rRNA read files were processed with DADA2^15^. Taxonomy was assigned to merged non-chimeric amplicon sequence variants (ASVs) using the SILVA rRNA database (version 138.1)^16^. Potential contaminating ASVs were filtered with Decontam using the prevalence method^17^. Three types of negative controls were included: sampling controls from the clinic environment, DNA extraction blanks, and 16S rRNA PCR amplification blanks. Bioinformatic decontamination was benchmarked on serial dilutions of the mock microbial community standard ^18^. Separate sequencing runs were denoted as batches and accounted for in all analyses. Differential abundance testing was conducted with MaAsLin2^19^.

### 2.6 Bacterial Whole Genome Sequencing and Bioinformatic Analysis

Preliminary colony identification was obtained by plating on CHROMagar™ Orientation media (RT412). Prior to whole genome sequencing, species-level colony identification was performed by MALDI-TOF (VITEK® MS PRIME, bioMérieux). Genomic DNA was isolated from a single source colony using the PureLink™ Genomic DNA Mini Kit (K182002, Invitrogen). Sequencing libraries were prepared with Illumina TruSeq adapters and sequenced on an Illumina NovaSeq 6000 or NovaSeqX at Vanderbilt Technologies for Advanced Genomics (VANTAGE) aiming for a minimum of 150x coverage (**Supplementary Table 3**). Isolates BTF6T1, BTF15T-1, BTF28T-2, BTF39T-2, and BTF39T-1 were also sequenced using Oxford Nanopore technology at Plasmidsaurus Inc.

The genomic definition of a bacterial *strain* remains under debate ^20–23^. Informed by prior research, we considered several approaches to determining what boundary constitutes a bacterial *strain*, including average nucleotide identity (ANI) and core genome single nucleotide variants (SNVs). For example, Thänert *et al*. utilized a pairwise core genome SNV threshold of 500 to determine uropathogenic *E. coli* (UPEC) “lineages” for evolutionary comparison^24^. A similarly designed study of within-patient *Salmonella* evolution considered strains as those with fewer than 36 pairwise SNVs compared to a reference genome^25^. ANI thresholds of 99.95% or 99.999% have also been employed to define bacterial strain boundaries, particularly strains within metagenomic samples^25–28^. We calculated ANI with fastANI (version 1.3) using a minimum fragment length of 3,000 bp^29^. To evaluate core genome SNVs of *E. coli* genomes, we utilized a collection of previously published urinary *E. coli* isolates from our laboratory; each isolate was collected from a unique patient (NCBI BioProject PRJNA819016)^30–33^. We created a core genome alignment with panaroo (version 1.4.1) and determined pairwise SNVs with snp-dists (version 0.8.2)^34,35^. We observed concordance between thresholds of >99.99% ANI and <100 core genome SNVs (**Extended Data Fig. 1**). All genomes with >99.99% also had core genome SNVs <100. Several pairs of genomes from separate patients had <100 core genome SNVs but less than 99.99% ANI (**Extended Data Fig. 1b**). We therefore used thresholds of >99.99% ANI and <100 pairwise core genome SNVs to define bacterial strains in this study. These thresholds are also intuitive in the context of previously reported mutations rates of *E. coli* (1-10 SNVs per genome per year) and the time scale separating the isolates in this present study (**Supplementary Table 7**)^36–41^.

The presence of antimicrobial resistance genes (ARGs) and stress response genes was predicted from assembled genomes using Resistance Gene Identifier (RGI, version 6.0.3) which implements the Comprehensive Antibiotic Resistance Database (CARD, version 4.0.0)^42^. Additionally, ARGs were predicted with ResFinder (version 4.6.0) as an orthogonal approach^43,44^. Chromosomal point mutations for *E. coli* and *Klebsiella* spp. were predicted with taxa-specific databases with PointFinder (version 4.1.1)^45^. ARGs are detailed in **Supplementary Table 5**. Plasmids content was predicted from genome assemblies using the MOB-suite (version 3.1.9)^46,47^. Predicted plasmids, replicon types, relaxase type, and predicted plasmid mobility are listed in **Supplementary Table 6.**

For identification of mutations within the isolates from the same patient, we used breseq (version 0.35.0) to map trimmed reads to the oldest isolate of a given patient’s strain (“reference isolate”, **Supplementary Table 7**)^48,49^. Genbank annotation files produced by Bakta were used as the reference genomes for breseq read mapping. Breseq was run in consensus mode with a frequency cutoff of 0.8 to identify mutations, including single nucleotide variants (SNVs), insertions, deletions, and structural variations. Putative variants were manually inspected to exclude artifacts based on coverage depth, mutation frequency, and quality scores. Variants are detailed in **Supplementary Table 8**. Complete breseq output is uploaded to the accompanying GitHub page.

For gene ontology (GO) over-representation analysis, we re-formatted breseq output with gdtools and extracted gene locus tags from nonsynonymous and nonsense mutations, and from insertions and deletions in coding sequences. We extracted gene ontology (GO) terms from the Bakta annotation file based on the gene locus tag. Each gene may be annotated with one or more GO terms corresponding to the three GO stratifications: biological process, molecular function, and cellular component. We compared the frequency of GO terms in the mutation dataset to the frequency of GO in the entire genome via a Fisher’s exact test with multiple testing correction with the Benjamini-Hochberg procedure (**Supplementary Table 9**). Corrected P values are reported as false discovery rates (FDR).

### 2.7 Genetic Manipulation of LPS/O-antigen Biosynthesis Genes in *E. coli*

Bacterial strains and plasmids used for genetic manipulation are listed in **Supplementary Table 10**. Bacterial strains were grown in lysogeny broth (LB) at 37°C. For selection during cloning, ampicillin (100 µg/mL) and kanamycin (50 µg/mL) were used. Deletion of the *waaW* (UTI89_C4169) and *waaL* (UTI89_C4167) genes was performed in the model cystitis strain UTI89 using the λ Red recombinase system of Murphy and Campellone^50–54^. Electrocompetent cells were prepared using a 20% glycerol and 0.1mM MOPS buffer. Electroporation was conducted using the Gene-Pulser instrument (1.5 kV, 400 Ω resistance, and 25 µF capacitance, Bio-Rad). Successful deletions were confirmed by whole genome sequencing (UTI89Δ*waaW*, BioSample SAMN44467476) or by colony PCR (**Supplementary Table 11**). The complementation construct for *waaW* (pWaaW) was generated in the pTRC99a plasmid backbone with *waaW* cloned under its native promoter. Briefly, the *waaW* gene was amplified from genomic DNA, whereas the pTRC99a vector was digested with EcoRI and KnpI. The *waaW* amplicon and digested vector were assembled by Gibson Assembly (E5510S). Complemented strains were grown in LB with 100 µg/mL ampicillin. Correct plasmid assembly was verified by Oxford Nanopore sequencing at Plasmidsaurus Inc. Oligonucleotide primers for cloning are listed in **Supplementary Table 11**.

### 2.8 LPS Extraction and Staining

LPS was extracted from bacterial cultures using successive aqueous-phenol extractions^55^. Briefly, 0.5mL of an overnight culture was centrifuged, and the pellet was resuspended in Laemmli sample buffer (4% 2-mercaptoethanol, 4% SDS and 20% glycerol in 0.1 M Tris-HCl, pH 6.8, bromophenol blue). The suspension was treated with Proteinase K (NEB, P8107S) for 3 hours at 60 °C. Then, two successive phenol-ether extractions were performed as described by Davis and Goldberg^55^. Extracted LPS was separated by SDS-PAGE and silver stained with modifications previously described^56,57^. An LPS standard from *E. coli* O55:B5 was included alongside samples from this study (1.25 µg, Sigma-Aldrich, L5418). Following electrophoresis, the gel was briefly washed with Milli-Q H_2_O then fixed and oxidized in a single 15-minute step (40% v/v ethanol, 5% v/v acetic acid, 1% w/v periodic acid). The gel was washed 3x with Milli-Q H_2_O, then stained and developed with a commercially available silver stain kit (Pierce, 24612). The gel was visualized on a Gel Doc EZ System instrument (Bio-Rad).

### 2.9 Bacterial Adherence to Bladder Epithelial Cells

Bladder epithelial cells (ATCC 5637, HTB9) were maintained in RPMI 1640 Media (Gibco A1049101) with 10% heat inactivated fetal bovine serum (FBS), 100 IU/mL of penicillin and 100 μg/mL of streptomycin. Bladder epithelial cells were incubated at 37°C with 5% CO_2_. Bladder epithelial cell monolayers were infected according to previously published protocols^58,59^. Bladder epithelial cells were seeded at a density of 1×10^5^ cells per well of 24-well tissue culture-treated plates (Fisher/Falcon, 087721H) ∼18 hours prior to infection in media without antibiotics. UTI89 and its isogenic mutants were cultured statically for ∼24 hours in LB at 37°C, followed by a 1:1000 sub-culture into fresh LB and an additional 24 hours of static growth to induce type-1 pili production^53^. Cultures were normalized to an OD_600_ 3.4 (∼2×10^9^ CFUs/mL) in PBS. Bladder epithelial cells were infected at a multiplicity of infection (MOI) of 5-10 CFUs. Following infection, 24-well plates were centrifuged at 600×g for 5 minutes to synchronize bacterial adherence. Then, bladder epithelial cells were incubated for 2 hours to allow bacterial interactions with the epithelial cells. Next, one set of wells was lysed with 0.1% v/v Triton X100 added directly to the RPMI to determine the total CFUs per well. Another set of wells was washed 4x with PBS to remove planktonic or loosely adherent bacteria and then lysed with 0.1% v/v Triton X100. Finally, a set of wells was washed with PBS and then incubated with the eukaryotic impermeable antibiotic gentamicin (100 µg/mL) for 2 hours to kill extracellular bacteria. Following 2 hours of antibiotic treatment, the wells were washed with PBS and lysed with 0.1% v/v Triton X100. Following lysis of bladder epithelial cells, samples were serially diluted in PBS and plated on LB-agar to enumerate CFUs.

### 2.10 Macrophage Internalization and Survival Assay

Macrophage infection assays were slightly modified from previously published protocols^60,61^. Immortalized murine macrophages (RAW 264.7) were cultured in RPMI 1640 media (ATCC modification) with 10% v/v FBS in 5% CO_2_ atmosphere at 37°C. Macrophages were seeded to a density of 1×10^5^ cells/well in a 24-well tissue culture plate 18-hours prior to infection. Then, bacteria suspended in PBS were added to each well at an MOI of 10. The 24-well plate was centrifuged at 600g for 5 min to expedite bacterial contact with the macrophages. Bacterial internalization was allowed to proceed for 30 min at 37 °C. One set of wells was lysed with 0.1% v/v Triton X100 (ThermoFisher, A16046AE); this represents the total CFUs in each well. Another set of wells was washed 3x with PBS and then lysed with 0.1% Triton X100; this represents internalized bacteria at 30 minutes post-infection. A final set of wells were treated with RPMI + 100 µg/mL of gentamicin (Gibco, 15750060) to kill extracellular bacteria, incubated for 1 hour to allow macrophage killing of internalized bacteria, washed with PBS 3x, and then lysed with 0.1% Triton X100. Following lysis of macrophages, the lysate was serially diluted in PBS and plated on LB-agar plates to enumerate CFUs.

### 2.11 Mouse Infections

Mouse infection protocols were reviewed and approved by the Vanderbilt University Medical Center Institutional Animal Care and Use Committee (IACUC) (protocol # M1800101-01). Mouse infections were conducted in 5–8-week-old C3H/HeNHsd female mice (Envigo/Inotiv Inc.) following previously published methods ^62,63^. Strain UTI89 and its isogenic mutants were cultured statically for ∼24 hours in LB at 37°C, followed by a 1:1000 sub-culture into fresh LB and an additional 24 hours of static growth to induce type-1 pili production^53^. To differentiate inoculated strains from commensal *E. coli* harbored in Envigo mice, we utilized the strains containing the kanamycin-resistance cassette in place of the gene of interest (i.e., marked deletion), and all samples were plated on LB agar supplemented with kanamycin at a final concentration of 50 µg/mL. Fecal pellets were collected in pre-weighed tubes containing 1mL of PBS. Mice were anesthetized with 3% isoflurane and transurethrally inoculated with ∼10^7^ CFUs resuspended in 50µL of PBS. Mice were humanely euthanized at either 24- or 72-hours post-infection. Organs were aseptically dissected and homogenized in sterile PBS. Homogenates and fecal suspensions were serially diluted in PBS and plated on LB-agar to count CFUs.

### 2.12 Bacterial Motility Assays

Swimming motility was measured in soft LB agar (0.25% w/v) supplemented with triphenyl tetrazolium chloride (0.002% w/v). Assays were conducted in 6-well culture plates (Fisher/Falcon 0877249). Overnight cultures were inoculated into the agar in the center of each well with an inoculating needle. After 8 hours of incubation at 37°C, the diameters of bacterial spread were measured in millimeters. Experiments were completed 3 times independently. Technical replicate measurements were made perpendicular to the first measurement.

### 2.13 Phage Susceptibility Testing Methods

Bacterial strains were grown overnight in LB at 37 °C. 70 μL of overnight culture was mixed with 6 mL of molten 0.5% agar LB and poured onto (Thermo Scientific NUNC OmniTray) plates with a bottom layer of 1.2% agar LB. Phage titers were serially diluted 10-fold in SM buffer (H_2_O, 50mM Tris-HCl pH 7.4, 5.8 g/L NaCl and 8mM MgSO_4_) and 2 μL of each well was spotted onto the plates and left to grow overnight at 37 °C. Plates were imaged the following day and efficiency of plaquing (EOP) ratios were calculated by comparing the number of plaques formed on the mutant strain compared to the parental strain on a given day. The spotting assays were conducted twice, and the values reported are the average between the two.

## 3. RESULTS

### 3.1 Study Cohort and Microbiome Characteristics of Children with Spina Bifida

We collected urine and fecal samples from 77 children and young adults with spina bifida from two medical centers (**Fig. 1**). Hand swabs were collected from patients who catheterized (**Fig. 1b**). The age range of our cohort was 0 – 27.4 years old (**Fig. 1c and Supplementary Table 1**). The cohort was composed of 36 females and 41 males. Forty-four subjects (57.1%) performed clean intermittent catheterization (CIC) for bladder management. Twenty-three of the subjects (29.9%) had received antibiotics in the past 30 days, and forty-nine subjects (63.6%) had a history of UTI (**Fig. 1c and Supplementary Table 1**).

**Figure 1.**
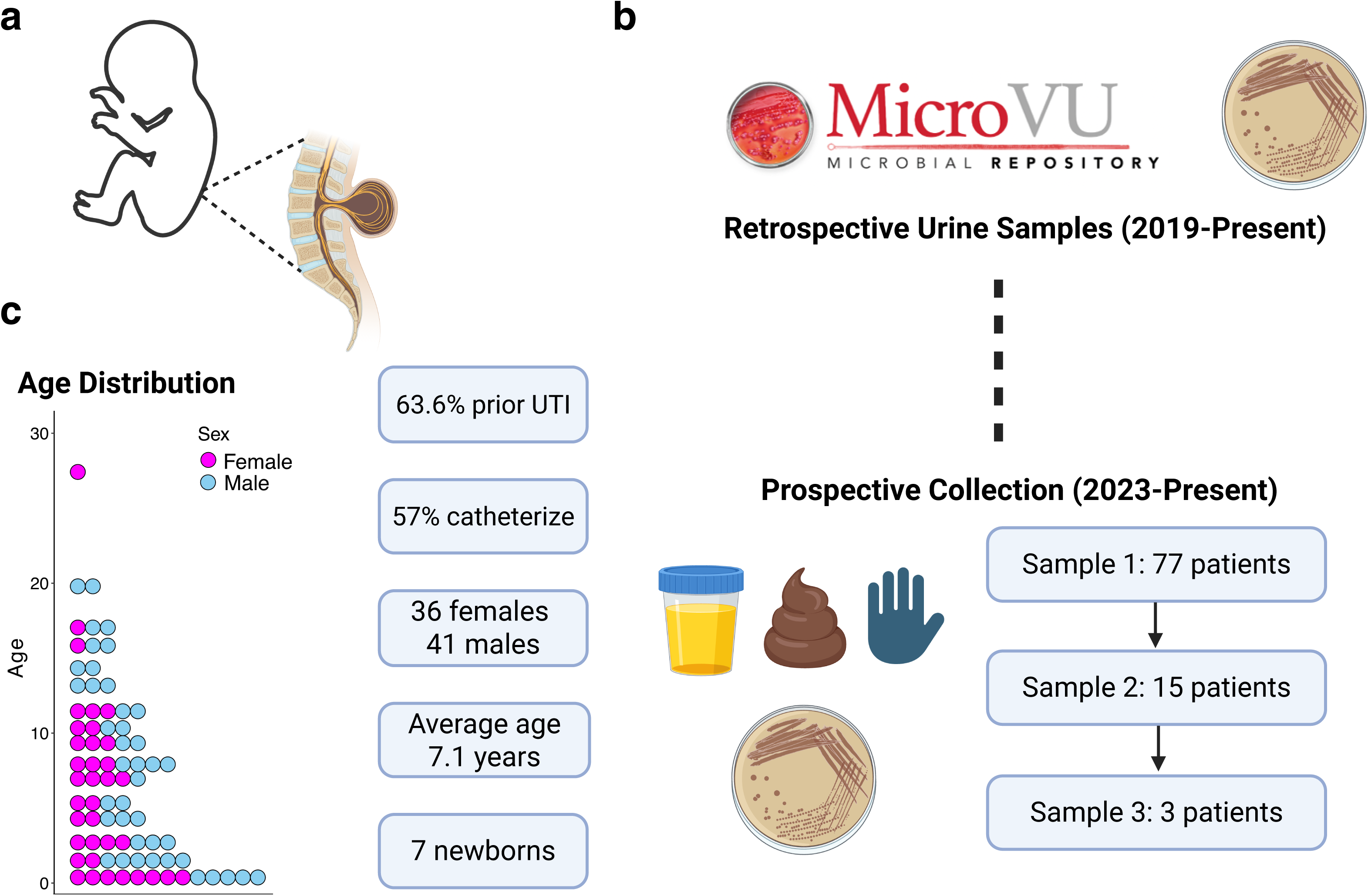
Study schematic. **a,** Schematic representation of spina bifida, a congenital malformation of the spina cord. Spina bifida leads to neurogenic bladder and frequent urinary tract infections (UTIs). **b,** Study protocol overview. Prospective sample collection of urine and fecal samples, and hand swabs began in the year 2023 at two medical centers. Bacterial isolates from historical urine samples from enrolled patients were accessed in the microbial biorepository MicroVU at Vanderbilt University Medical Center. **c,** Age distribution in 1 year bins and key cohort metrics. Each dot represents an individual patient. Additional cohort metrics are detailed in Supplementary Table 1. Figure created with BioRender under an institutional license.

We performed 16S rRNA amplicon sequencing on urine and fecal samples, and hand swabs (**Fig. 2a**). Samples were collected from patients without signs or symptoms of UTI at the time of collection. In urine samples, we noted a high abundance of taxonomic families representative of urinary pathogens (e.g. *Enterobacteriaceae* and *Enterococcaceae*) and the absence of health-associated taxa such as *Lactobacillaceae* and *Prevotellaceae* (**Fig. 2a**). In general, urine samples exhibited low alpha diversity and were often comprised of a single taxonomic family.

**Figure 2.**
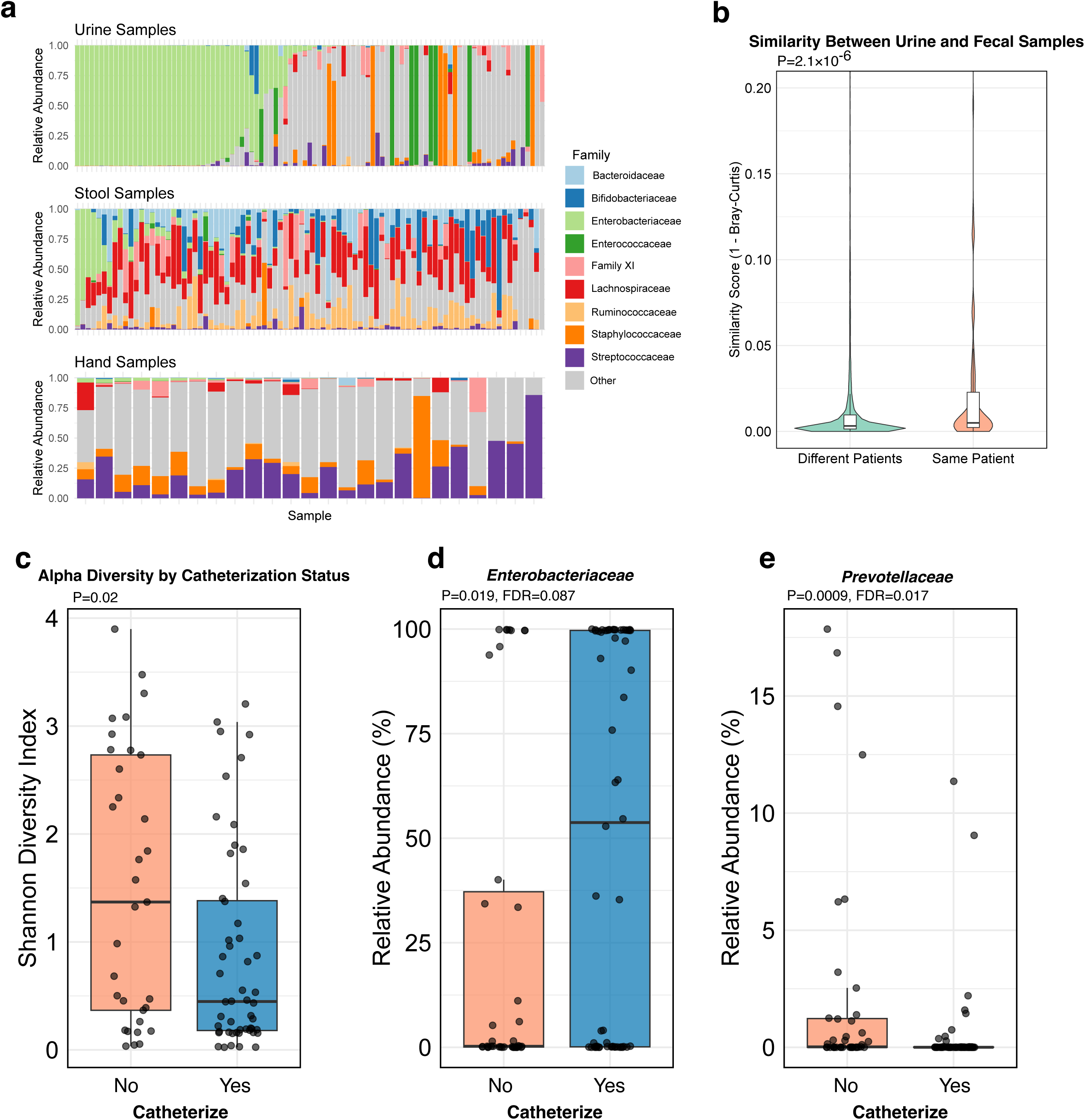
Microbiome characteristics of urine, fecal, and hand swab samples from children with spina bifida. **a,** Stacked bar plots of bacterial taxonomic family relative abundance for urine, fecal, and hand swab samples. **b,** Microbiome similarity between urine and fecal samples from the same or different patients. Microbiome similarity was calculated as 1-Bray-Curtis dissimilarity distance. P value calculated with Wilcoxon rank sum test. **c,** Shannon diversity of urine samples stratified by catheterization status. P value calculated with Wilcoxon rank sum test. **d-e,** Relative abundance of taxonomic families *Enterobacteriaceae* and *Prevotellaceae* stratified by catheterization status. Differential abundance testing was conducted with MaAsLin2, correcting for age, sex, sequencing batch as fixed effects in the linear model, and accounting for repeated measures by including subject ID as a random effect. Benjamini– Hochberg corrected P values are reported as false discovery rates (FDR).

We hypothesized that manipulation of the urinary tract or catheterization increases the potential of cross-contamination between urinary and fecal microbiota, facilitating microbial transfer between the two ecosystems. We computed the Bray-Curtis distance between urine and fecal samples and plotted a similarity score (1-Bray Curtis Distance) for urine-fecal pairs from the same or different patients (**Fig. 2b**). Indeed, microbiome similarity was significantly higher *within* patients compared to *between* patients (**Fig. 2b**, Wilcoxon P=2.1×10^-6^), supporting the hypothesis of microbial exchange between urine and fecal microbiomes. We compared 16S rRNA amplicon sequence variants (ASVs) between urine and fecal samples. We focused on urine samples containing *Escherichia*, *Enterococcus*, or *Klebsiella* ASVs.

Eighty-one (81) of 97 urine samples (83.5%) had *Escherichia* ASVs, 50 urine samples (51.5%) had *Enterococcus* ASVs, whereas 12 urine samples (12.4%) had *Klebsiella* ASVs. Of the 81 urine samples with *Escherichia* ASVs, 67 paired fecal samples (82.7%) shared the same *Escherichia* ASV or ASVs. In contrast, only 1 fecal sample shared *Klebsiella* ASVs with a paired urine sample (8.3%). Twenty-seven (27) fecal samples shared *Enterococcus* ASVs with paired urine samples also containing *Enterococcus* ASVs (54%). Many patients with neurogenic bladder must catheterize to prevent urinary retention. We compared urine samples from children who catheterize from those who do not catheterize. Among patients who catheterized, we noted decreased urine alpha-diversity (**Fig. 2c**). Moreover, patients who catheterized has significantly increased *Enterobacteriaceae* and reduced *Prevotellaceae* (**Fig. 2d-e**). Together, these data demonstrate concordance between urine and fecal samples in children with spina bifida and highlight how common urologic interventions shape microbiome diversity and composition.

### 3.2 Whole Genome Sequencing of Longitudinally Collected Bacterial Isolates

Based on our observation that urine samples were frequently dominated by urinary pathogens such as *E. coli*, *Enterococcus* spp., and *Klebsiella* spp., we next asked whether the same bacterial strains were present within individual patients across time. We performed whole genome sequencing on cultured bacterial isolates collected during this study or retrieved from the MicroVU microbial biobank. Isolates from MicroVU allowed us to sample historical isolates from the same patients beginning in 2019. We sequenced bacterial isolates from patients for whom multiple isolates were available for comparison. We sequenced 97 isolates from 24 unique patients (**Supplementary Table 3**). The average number of isolates per patient was 3.75 (range 2-9 isolates) and an average of 3.1 timepoints (range 2-8 timepoints). The patients included in the whole genome sequencing spanned ages 3 months to 19 years old.

We identified two predominant patterns of urinary tract colonization (**Fig. 3**). We observed the rapid cycling of strains and/or species, often following antibiotic treatment (e.g., BTF10 & BTF61). We also observed the persistence of a single strain within a patient across timepoints (denoted by asterisks in **Fig. 3**). We defined strains as isolates from the same species with >99.99% average nucleotide identity and fewer than 100 core genome SNVs between isolates (**Methods**). No bacterial genomes from separate patients exceeded 99.99% ANI, indicating the absence of transmission between patients. The persistence of a single strain within a patient occurred in ∼33% of the strains sequenced (32/97 isolates sequenced had a paired strain). This pattern diverges from the pattern observed in otherwise healthy individuals with recurrent UTI (rUTI) where a single strain is implicated in *most* recurrent infections^24,64–68^. The presence of an identical strain across timepoints allowed us to investigate in-patient pathogen evolutionary changes to understand how the host environment may be shaping persistence in these isolates.

**Figure 3.**
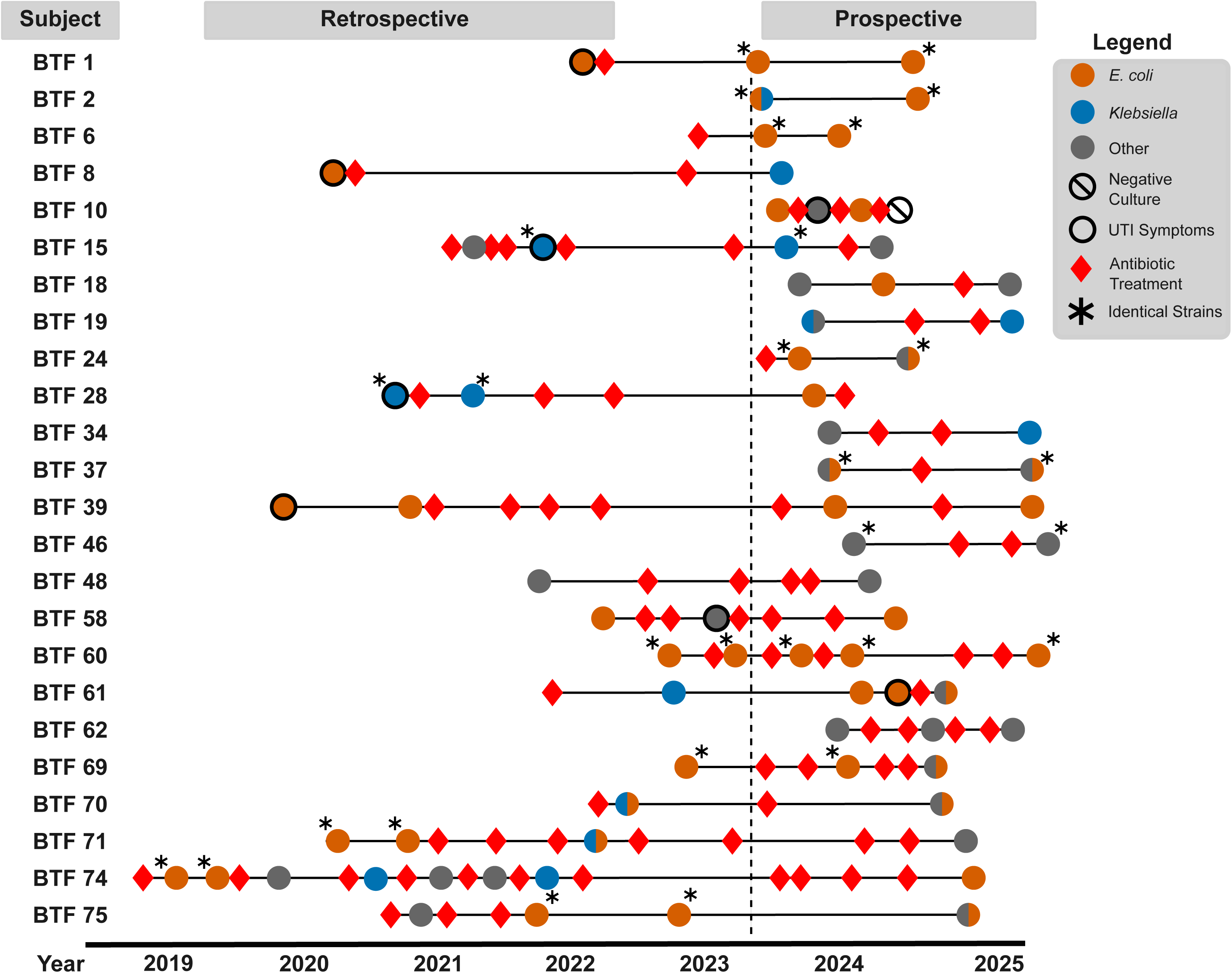
Longitudinal urinary isolates reveal patterns of ecological succession. Whole genome sequencing of bacterial isolates. Patients with multiple cultured isolates were included. Individual patients are represented on the left vertical axis. The vertical dashed line depicts when prospective sample collection began. The asterisks denote strains that were present *within* the same patient at multiple timepoints (**Supplementary Table 7**). No strains were shared *between* patients. Retrospective samples were accessed from the MicroVU microbial biorepository at Vanderbilt University Medical Center. Only bacterial isolates available for sequencing are depicted. Antibiotic usage and symptomatology were extracted from medical record chart review. The location of symbols on the schematic is approximate.

First, we measured phenotypic antibiotic susceptibility of all sequenced bacterial isolates using broth microdilution assays (**Supplementary Table 4**). In parallel, we predicted antimicrobial resistance genes (ARGs) within genome assemblies with ResFinder and the Comprehensive Antibiotic Resistance Database (CARD) (**Supplementary Table 5**). ARG content varied widely based on bacterial taxa (**Extended Data Fig. 2**). We compared the ARG content between successive isolates from the same patient. We considered 3 scenarios (**Fig. 3**): the same strain at successive timepoints, the same species at successive timepoints, and the different species at successive timepoints. ARG content remained similar between successive isolates of the same strain or same species; that is, there was no net gain or loss of ARGs between successive strains (**Extended Data Fig. 3**). The ARG content between successive bacterial isolates of different species varied widely, based on the intrinsic differences in ARG repertoires between taxa, yet remained centered around zero net change in ARG content (**Extended Data Fig. 3**). Likewise, phenotypic susceptibility to antibiotics did not change dramatically between successive isolates of the same strain (**Supplementary Table 4**). This indicates that persistence of a single strain is not facilitated by increasing antibiotic resistance. Likewise, recolonization by a new strain or species does not consistently trend towards more resistant-organisms over the time scale that we sampled. Similarly, plasmid content remained relatively stable between timepoints of the same strain (**Extended Data Fig. 4**). These observations suggest that, in the sub-cohort of patients with whole genome sequencing results, acquisition of antibiotic resistance is not the primary driver of bacterial persistence in or re-colonization of the urinary tract.

### 3.3 Within-Patient Urinary Pathogen Evolution

In order to begin to elucidate molecular factors that may contribute to longitudinal carriage of a clonal strain within a patient, we took advantage of the 15 instances in which a patient was colonized with the identical strain across timepoints (denoted by asterisks in **Fig. 3**). In our collection we observed examples of longitudinal carriage of multiple urinary pathogen species including 11 *E. coli* strains (**Fig. 4a**), two *Klebsiella pneumoniae* strains, one *Proteus mirabilis* strain, and one *Enterococcus faecalis* strain (**Supplementary Table 7**). The intervals between which these isolates were collected spanned 46 days to 811 days (**Supplementary Table 7**). Multiple instances of shared *E. coli* strains with individual patients allowed us to estimate a molecular clock of *E. coli* evolution within patients. We extracted core genome SNVs between sequential isolates from a core genome alignment and performed linear regression (**Fig. 4b**). The slope was 0.076 core genome SNVs per day or 27.7 core genome SNVs per year (P=0.02, 95% confidence interval 0.01157 to 0.1399 core genome SNVs per day). This estimate was elevated by isolate BTF75T1 which contained 73 core genome SNVs relative to fewer than 20 core genome SNVs in all other *E. coli* strain pairs.

**Figure 4.**
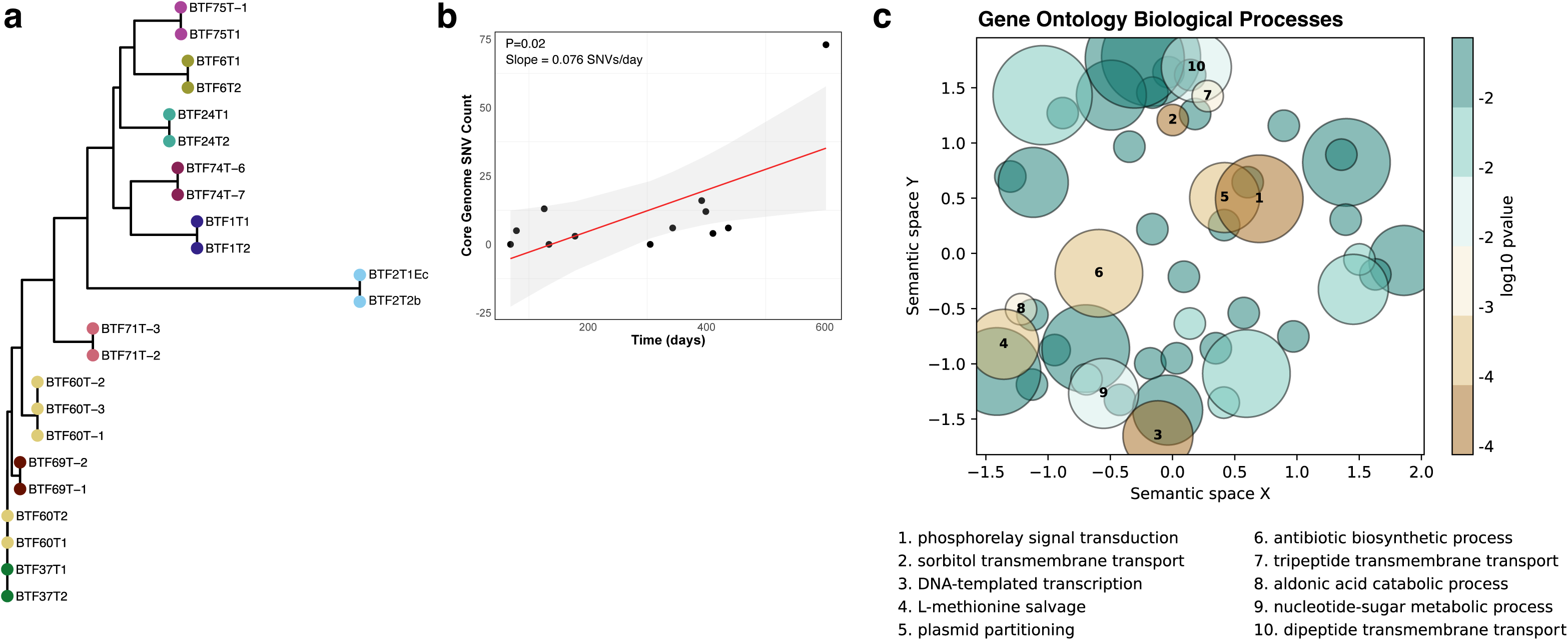
Bacterial whole genome sequencing reveals mutational patterns in longitudinal isolates. **a,** Phylogeny of *E. coli* isolates from shared strains within individual patients (**Supplementary Table 7**). Node colors represent unique patients. The phylogenetic tree was generated with FastTree and plotted with ggtree^103^. **b,** Core genome SNVs vs. time (days). Core genome SNVs were extracted from a panaroo core genome alignment with snp-dists and plotted in R. **c,** Biological process GO terms were clustered by semantic similarity with GO-Figure!^104^. The size of the bubble is proportional to the number of genes associated with the particular semantic cluster. The color represents the Fisher exact test P-value from GO overrepresentation analysis (**Supplementary Table 9**).

We used breseq to map reads to each strain’s oldest isolate (“reference isolate” in **Supplementary Table 7**). We aggregated mutations across patient isolates to highlight patterns of evolution across patients and colonizing urinary pathogen species (**Supplementary Table 8**). We focused on non-synonymous mutations and nonsense mutations, and deletions and insertions in coding sequences since these mutation types are most likely to impact protein function. Mutations were predominantly found in genes encoding key components of the bacterial cell envelope, particularly those involved in immune or phage evasion (**Table 1**). These included genes for iron acquisition systems, various cell envelope and flagellar proteins, and components of the LPS/O-antigen biosynthesis pathway. Notably, these categories of genes were consistently mutated across multiple bacterial species, indicating that this pattern is not unique to *E. coli* (**Table 1**).

**Table 1.**
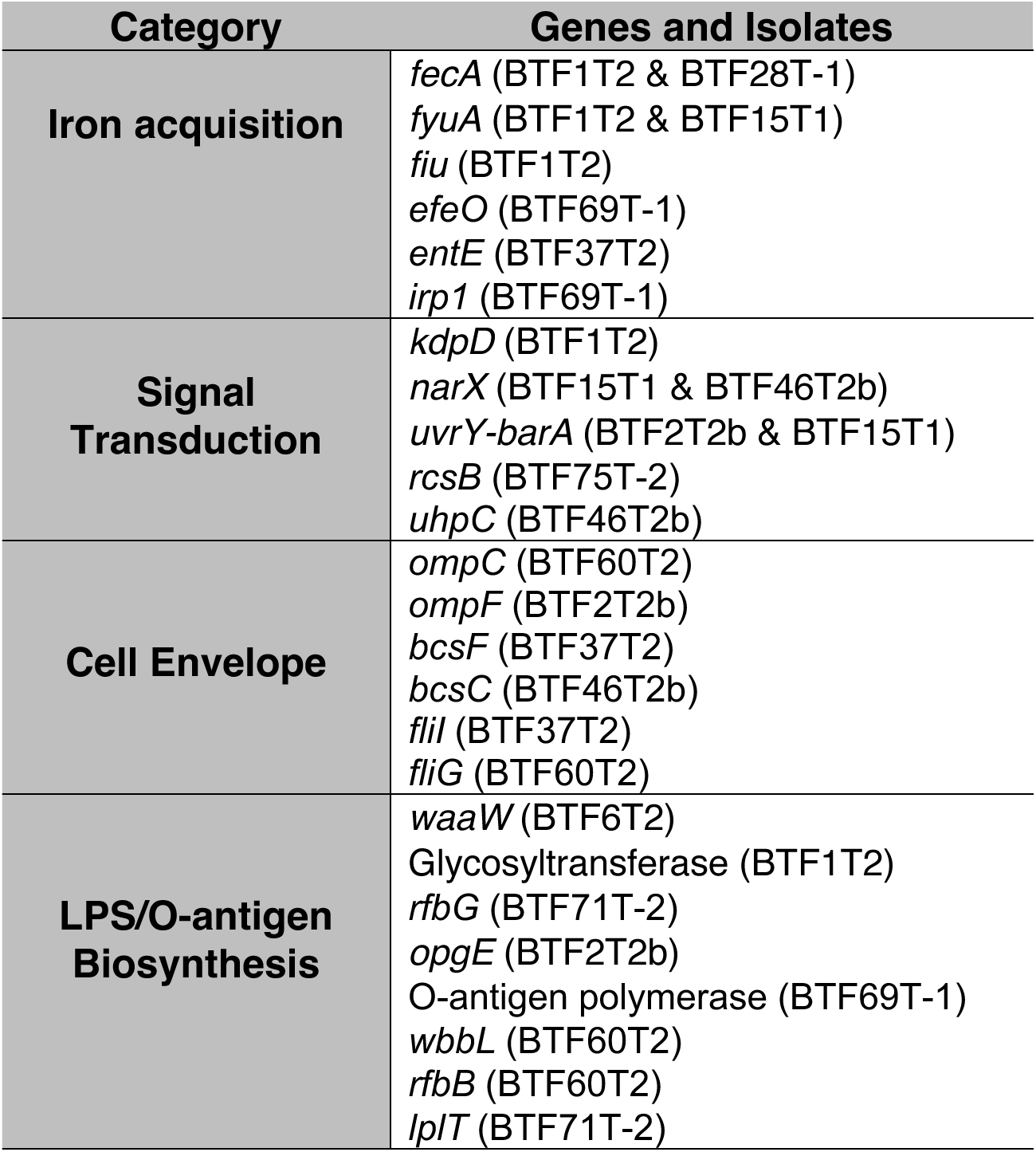
Functional Groups of Mutated Genes.

We sought to determine whether non-synonymous and nonsense mutations, and deletions and insertions in coding sequences occurred disproportionately in any particular gene ontology (GO) categories. The null hypothesis—representing neutral selection—is that mutations should occur proportionally to the percent of the genome that those specific gene ontologies occupy. We compared the frequency of GO terms in the mutations among longitudinal isolates to the frequency of those GO terms in the entire genome annotations of the longitudinal isolates. There were 276 unique GO terms among the genes with mutations (**Supplementary Table 9**). The GO classifications overrepresented in the mutation dataset included GO terms related to mobile genetic elements including transposases and plasmids (GO:0004803, GO:0030541, FDR<0.05; and GO:0006276, FDR<0.1). Consistent with multiple mutations in two component signaling systems (TCSs) (**Table 1**), we observed overrepresentation of GO terms related to signaling receptor activity (GO:0038023, GO:0000160, FDR<0.05) and transcription (GO:0006351, GO:0003677, FDR<0.05) (**Fig. 4c**). An additional cluster of overrepresented GO terms was comprised of small molecule transport ontologies including dipeptide and tripeptide transport (GO:0035442 and GO:0035443, respectively, both FDR<0.05), siderophore uptake (GO:0015344, FDR<0.05), and sorbitol transport (GO:0015795, FDR<0.05) (**Fig. 4c**). The overrepresentation of sorbitol transport was driven by the deletion of the entire *sgc* operon in isolate BTF24T2 (**Supplementary Table 8**). GO terms related to the bacterial cell membrane were not significantly overrepresented in the mutational dataset relative to the background frequencies of such GO terms in the genomes (**Extended Data Fig. 5, Supplementary Table 9**). Genes related to LPS/O-antigen biosynthesis genes were not consistently annotated with GO terms, explaining their absence in the GO overrepresentation dataset. Based on the repeated patterns of gene categories overrepresented in mutations from longitudinal strains, we hypothesized that these mutations may correspond to increased bacterial fitness during long-term bladder colonization.

Another notable category of mutated genes included two-component signal transduction systems—bacterial-specific proteins that are absent in higher eukaryotes and play a crucial role in sensing and responding to environmental signals. We observed mutations in: *kpdD* (responsive to potassium limitation), *uvrY-barA* (responsive to short chain fatty acids), *rcsB* (responsive to osmotic stress), *uhpC* (regulates glucose-6-phosphate uptake), and *narX* (senses nitrate availability). Our group has previously systematically studied the role of TCSs in UTI pathogenesis and identified a role for the *narXL* TCS in bladder colonization^69^.

Another pattern of mutations was in genes involved in iron acquisition. Two isolates, one *E. coli* and one *K. pneumoniae* from patients BTF1 & BTF15, respectively, had mutations in the yersiniabactin siderophore receptor *fyuA* relative to the prior isolate of the same strain (**Table 1**). Isolate BTF1T2 had an additional mutation in *fiu,* a catecholate siderophore receptor. Likewise, two independent mutations in *fecA*, a ferric citrate transporter, were observed in one *E. coli* and one *K. pneumoniae* isolates (BTF1T2 & BTF28T-1, respectively). These mutations could suggest changes in iron-acquisition or alternative fitness advantages such as decreased phage susceptibility as siderophore receptors serve as targets for phage^70^.

Additionally, mutations in outer membrane porins such as *ompC* and *ompF*, as well as in genes involved in cellulose and flagellar biosynthesis, can alter membrane permeability, biofilm formation, and motility (**Table 1**). Similarly, we detected multiple mutations in LPS/O-antigen biosynthesis genes, including multiple loss of function mutations (**Table 1** and **Supplementary Table 8**). Mutations in LPS and O-antigen biosynthesis genes can alter bacterial surface structures, potentially impacting immune evasion, phage susceptibility, and colonization efficiency. We hypothesized that these mutations may increase bacterial fitness during long-term bladder colonization.

### 3.4 Modification of LPS Facilitates Immune Evasion Without Tradeoffs in a Murine UTI Model

We selected one mutational class to characterize further. Two *E. coli* strains were collected 3 months apart from a single patient (patient ID BTF6). The patient did not receive antibiotics during this period and was asymptomatically colonized (**Fig. 3**). At timepoint 2 (isolate BTF6T2), a SNV (865 C>T) introduced a premature stop codon in *waaW* (Gln289Stop) (**Supplementary Table 8**). *WaaW* (previously *rfaJ*) encodes the *E. coli* UDP-galactose--(Galactosyl) LPS alpha-1,2-galactosyltransferase. WaaW adds a second galactose residue to the oligosaccharide outer core of LPS (**Fig. 5a**) ^71–73^. The mutation observed in BTF6T2 occurs in the glycosyltransferase family C-terminal domain, the enzymatic domain of WaaW, likely rendering the protein nonfunctional (**Fig. 5b).** Loss of WaaW function has been reported to prevent subsequent extension of the O-antigen^72,74–76^. To evaluate the functional consequences of this mutation, we next created a clean deletion of *waaW* in the model UPEC strain UTI89 ^53,54^. The UTI89 *waaW* (UTI89_C4169) shared 99.4% nucleotide identity over 100% coverage of the BTF6T1 *waaW* and 100% amino acid identity. UTI89 and BTF6T1 have identical operon structure representing the R1-type oligosaccharide core (**Extended Data Fig. 6**). Extracted LPS was separated by SDS-PAGE and visualized by silver staining. BTF6T2 and UTI89Δ*waaW* exhibited truncated core oligosaccharide bands and reduced intensity of the high-molecular weight bands which correspond to O-antigen repeats (**Fig. 5c**). Complementation of *waaW* on a plasmid restored O-antigen and core oligosaccharide bands. These strains recapitulated the LPS banding pattern of a UTI89Δ*waaL* mutant, which lacks the O-antigen ligase to catalyze the addition of O-antigen to the oligosaccharide core (**Fig. 5c**).

**Figure 5.**
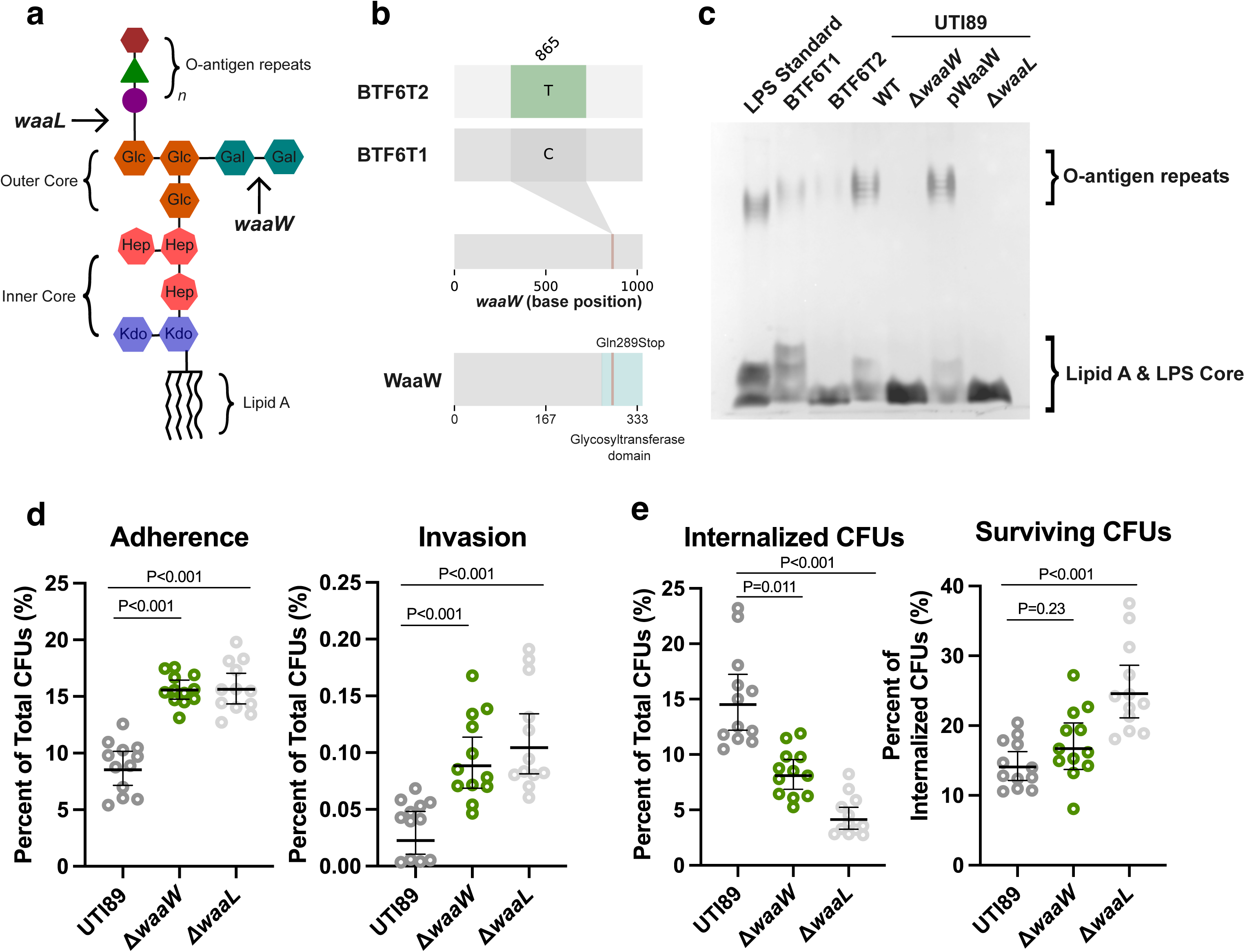
LPS/O-antigen biosynthesis mutations alter interactions with bladder epithelial cells and macrophages. **a,** Schematic of LPS with the R1-type core oligosaccharide. Geometric symbols represent sugar subunits of the oligosaccharide portions of LPS. Abbreviations: Kdo—3-deoxy-d-manno-oct-2-ulosonic acid; Hep—glycero-d-manno-heptose; Glc—glucose; Gal—galactose. **b,** Premature stop codon in *waaW* from *E. coli* isolate BTF6T2 relative to isolate BTF6T1 from the same patient. Mutation visualized with Snipit ^105^. **c,** Silver stained LPS extracts. The LPS standard--is a commercially available product from *E. coli* serotype O55:B5 (1.25 µg of LPS standard was added). **d,** Adherence to and invasion of bladder epithelial cells (ATCC HBT9). UTI89 and its isogenic mutants *waaW* and *waaL* were tested. **e,** Macrophage internalization and survival of UTI89 and its isogenic waaW mutant. Internalized CFUs were counted 30 minutes after inoculation, and surviving CFUs were counted from within the macrophages 90 minutes after inoculation. For panels **d** and **e**, the horizontal line is the geometric mean, and the error bars depict the 95% CI. P values were calculated with non-parametric Kruskal–Wallis test with two-sided Dunn’s *post hoc* test for multiple comparisons.

Previous work showed that swimming motility is impaired by mutations in O-antigen expression or processing^77^. We thus tested BTF6 clinical isolates and UTI89 clean deletion strains and complemented controls for their ability to swim in soft agar. While the clinical isolates BTF6T1 & BTF6T2 were both non-motile, deletion of *waaW* or *waaL* in UTI89 displayed impaired swimming motility (**Extended Data Fig. 7**), consistent with previous reports ^77^. Complementation of *waaW* in UTI89Δ*waaW* increased motility to nearly wild-type levels. Loss of swimming motility in LPS mutants has been previously attributed to cell envelope rearrangement and consequent loss of flagellar assembly^77,78^. This phenomenon has been described in the common laboratory *E. coli* K12 strain MG1655. Many MG1655 isolates contain a transposon inserted in *wbbL*, a rhamnose-transferase, ablating O-antigen production and rendering these strains non-motile^79–83^. Complementation of a functional *wbbL* restores O-antigen production and swimming motility in these strains^84,85^.

We hypothesized that the altered LPS structure may modulate *E. coli* interactions with host cells and modify the strain’s immunogenicity. We infected monolayers of bladder epithelial cells with UTI89, UTI89Δ*waaW*, or UTI89Δ*waaL*. Two hours after inoculation, we washed bladder epithelial cells with PBS and counted bacteria firmly adherent to the epithelial cells. Both UTI89Δ*waaW* and UTI89Δ*waaL* exhibited increased adherence to bladder epithelial cells compared to the wildtype strain (**Fig. 5d**). Likewise, after treating with gentamicin to kill extracellular bacteria, both UTI89Δ*waaW* and UTI89Δ*waaL* had higher intracellular bacterial counts (**Fig. 5d**). We surmise that this increased adherence and invasion may be attributed to greater exposure of bacterial adhesive fibers like type 1 pili due to the lack of O-antigen on the cell envelope.

To probe the role of O-antigen in *E. coli* interactions with phagocytic immune cells, we infected murine macrophages with UTI89, UTI89Δ*waaW*, or UTI89Δ*waaL*. After 30 minutes, fewer UTI89Δ*waaW* were internalized inside macrophages (**Fig. 5e**). After an additional 1-hour incubation, there were more surviving UTI89Δ*waaW* cells within the macrophages than wild-type UTI89 (**Fig. 5e**). These data indicate that the deletion of *waaW* decreases phagocytosis by macrophages and promotes survival within macrophages. Moreover, transurethral inoculation of mice with UTI89, UTI89Δ*waaW*, or UTI89Δ*waaL* demonstrated no fitness defect for O-antigen biosynthesis mutants in a mouse model of UTI at either 24- or 72-hours post-infection (**Extended Data Fig. 8**), suggesting no apparent fitness trade-offs imparted to

*E. coli* due to this cell envelope modification. Likewise, mutants lacking O-antigen, were able to migrate to the gut and displayed comparable gut colonization to the wild-type parent **(Extended Data Fig. 8c**), again suggesting no overt fitness trade-offs in the murine model tested herein.

O-antigen and membrane proteins (including siderophore receptors) are frequently targets of bacteriophages. It is therefore possible that changes in O-antigen and other cell envelope proteins may influence susceptibility to phage. Phage susceptibility is critical to investigate because: a) it could provide insights into the phage ecology in these specific patients that may be contributing to the selective pressures imposed on the pathogen, and b) investigating susceptibility of evolved strains to diverse phage may illuminate phage-driven therapeutic approaches tailored to the specific patient’s colonizing bacteria^86^. To begin to investigate this possibility, we performed phage plaque assays on selected strains with a subset of the BASEL phage collection^87,88^. We included *E. coli* strains UTI89, UTI89Δ*waaW*, or UTI89Δ*waaL* as well as two pairs of longitudinal samples with mutations in O-antigen biosynthesis genes and iron acquisition membrane proteins (subjects BTF1 and BTF6, **Table 1**). In general, clinical strains and UTI89 were more resistant to phage than the laboratory adapted *E. coli* K12 strain MG1655 (**Extended Data Fig. 9a-c**). Strain BTF1T2 was strikingly susceptible to Bas67 phage relative to its parental isolate BTF1T1 (**Extended Data Fig. 9a**); the terminal receptor of Bas67 is rough LPS which is likely to be more exposed due to a mutation in a glycosyltransferase in the O-antigen biosynthesis operon in strain BTF1T2^88^ (**Table 1**). Despite the *waaW* stop codon in isolate BTF6T2, this strain exhibited minimal differences in phage susceptibility compared to the parental isolate BTF6T1 (**Extended Data Fig. 9b**). Furthermore, phages (Bas31 and Bas67) unable to infect O-antigen expressing K12 *E. coli* (*wbbL*+)^87^ have increased plaquing on UTI89Δ*waaL* (**Extended Data Fig. 9c**). These data highlight strain-specific phage susceptibilities introduced by mutations in membrane proteins and LPS/O-antigen biosynthesis.

Together, this longitudinal evolutionary dataset displays mechanisms by which urinary pathogens adapt to the urinary tract and the vulnerabilities that these mutations introduce, potentially exposing targets for therapeutic intervention in specific host environments.

## 4. DISCUSSION

In this study, we established a large cohort of children and young adults with spina bifida to study the dynamics of the urobiome and evolution of urinary pathogens. We chose individuals with spina bifida because they have neurogenic bladder and have a variety of factors that predispose them to chronical colonization of their urinary tract with urinary pathogens. Distinguishing chronic colonization from active urinary infection is challenging in this group given their altered pelvic sensation, but it is critical given the life-threating risk for pyelonephritis and bacteremia. A greater understanding of the microbial ecosystem in spina bifida patients is likely to improve the diagnosis and treatment of UTIs in this patient population. We employed complementary methodologies of 16S rRNA sequencing and whole genome sequencing to uncover ecological dynamics in the urinary and intestinal microbiomes of spina bifida patients.

From whole genome sequencing data, we detected that patients were frequently re-colonized with different strains or species following antibiotic treatment (**Fig. 3**). Despite frequent antibiotic exposures in this patient cohort (**Fig. 3**), both phenotypic and genomically-predicted antibiotic resistance remained stable. Re-colonization was rapid following antibiotic treatment, suggesting constant re-inoculation of the urinary tract. We propose that ecological opportunism—that is, which strain of a urinary pathogen reaches the urinary tract first following antibiotics —rather than antibiotic resistance drives re-colonization. We believe these patterns may inform the understanding of urinary colonization and UTIs in other types of neurogenic bladder, particularly patients with spinal cord injuries. In a minority of samples, the same bacterial strain was present at multiple timepoints thus allowing evolutionary analysis. Multiple observations of mutations in specific gene ontologies—such as those involved in iron acquisition, membrane permeability, cellulose and flagellar biosynthesis, and signal transduction— suggests selective pressure acting on bacterial traits critical for persistence in the urinary environment (**Table 1**). For example, mutations in cell envelope genes and membrane transporters may modulate nutrient uptake or immune reactivity, while changes in cellulose and flagellar genes may reflect a trade-off between motility and biofilm formation. These frequently mutated gene functional groups highlight bacterial strategies for niche adaptation to the urinary tract. Given the cross-species mutations observed in this dataset, we propose that targeting such pathways could disrupt urinary pathogen colonization or persistence, offering promising avenues for therapeutic intervention in chronic bacterial colonization in neurogenic bladder. Likewise, the fitness costs induced by such mutations, such as decreased motility and phage susceptibility, highlight trade-offs of niche adaptation. These trade-offs may be exploited to develop new therapeutics or precision probiotics.

The role of O-antigen in UTI pathogenesis has been previously studied^89–94^. These prior studies suggested that loss of O-antigen reduced pathogenicity and persistence in mouse models of UTI^90,91^. In contrast, our data displays that loss of O-antigen facilitates immune evasion *in vitro* (**Fig. 5d-e**). This discrepancy with prior reports may be explained by relative antigenicity of different strains O-antigen type and/or the chronicity of infection. In a longitudinal study of patients with recurrent UTIs, positive selection of the rhamnosyltransferase-encoding gene *wbbL* was observed in both urinary and fecal *E. coli* isolates compared to prior isolates from the same patient’s UPEC strain^24^. Selection for and against O-antigen expression has previously been observed in *Burkholderia* spp. isolates from chronically infected cystic fibrosis patients^95–99^. In this context, O-antigen expression appears to be lost during early years of infection, and this loss may be advantageous for dissemination and transmission^97^. During later stages of chronic infection, the SNVs conferring loss of O-antigen revert to allow the production of O-antigen^96,97^. The expression of O-antigen may be a switch that modulates immunogenicity, dissemination potential, and infection chronicity. Furthermore, the loss of flagellar motility could indicate absent flagellar assembly^78^. Lack of flagella could impact infection dissemination or immunogenicity irrespective of the immune response induced by LPS.

Our ability to conduct strain-to-strain comparisons was limited by historically collected specimens and random biobanking of bacterial isolates before the conception of this present study. Thus, only a fraction of the bacterial isolates from the enrolled patients were available for historical comparison. More dense sampling intervals may allow additional comparisons between the same strains and further illuminate the dynamics of urinary pathogen colonization in spina bifida. Similarly, the sequencing of a single bacterial colony is unlikely to capture the vast diversity of a replicating bacterial population. This practice has been standard for bacterial evolution analysis^95–97,100,101^. A single colony may not illuminate low-frequency or emerging variants. There are well-studied examples of bacterial sub-populations exhibiting divergent phenotypes, such as antibiotic heteroresistance. Sequencing multiple or pooled colonies from the same urine sample may allow the detection of additional bacterial strains and evolution^28,102^.

## Supporting information

Gene ontology overrepresentation analysis.

Specimen metadata for 16S rRNA sequencing.

Cohort characteristics.

Bacterial strains and plasmids used for cloning.

Primers for cloning waa mutants.

Mutations from strains shared within individual patients.

Antimicrobial susceptibility of sequenced colonies.

Predicted plasmids from genome assemblies.

Antimicrobial resistance gene content of genome assemblies.

Strains shared within individual patients.

Bacterial genome accession and assembly metrics.

## DATA AVAILABILITY

Raw sequencing data is publicly available under NCBI BioProject PRJNA1155606. Bioinformatic code is available: https://github.com/reaset41/SpinaBifida-Urobiome.

## ACKNOWLEDGEMENTS

We thank the patients who participated in this study. This study was funded by the NIH under awards P20DK123967 (JES, DBC, and MH), R01AI168468 (MH), T32GM007347 (SAR), and F30AI169748 (SAR). The project was supported by Vanderbilt Institute for Clinical and Translational Research (VICTR) under CTSA award No. UL1 TR002243 from the National Center for Advancing Translational Sciences. MTL is an Investigator of the Howard Hughes Medical Institute. Graphics were created in Inkscape using open-source vector templates from SciDraw.io, bioart.niaid.nih.gov, bioicons.com, or with BioRender under an institutional license.

## AUTHOR CONTRIBUTIONS

SAR, BTF, DBC, and MH conceptualized the study. SAR, BTF, JES, MTL, DC, and MH acquired funding for this study. MTL, MSK, DBC, and MH supervised the study. AB, BTF, MSK, MKG, and SAR collected and processed patient samples. SAR and OFH processed, analyzed, and interpreted the sequencing data. SAR, ET, LRB, CBWS, and HG performed laboratory experiments. SAR and MH wrote the manuscript with input from all authors.

---

**Extended Data Figure 1.**
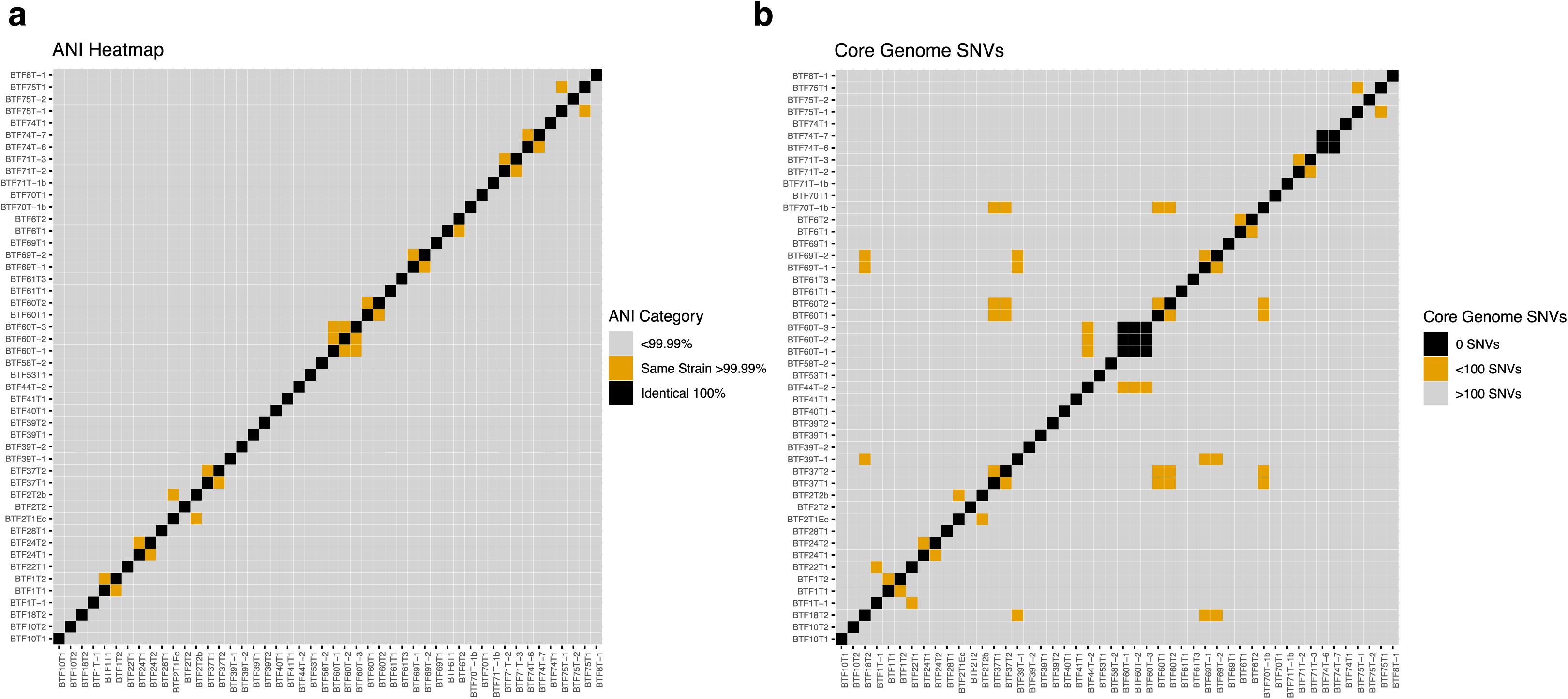
Benchmarking strain thresholds with average nucleotide identity (ANI) and core genome single nucleotide variants (SNVs). **a,** ANI of 48 *E. coli* genome assemblies from this study. ANI was calculated with fastANI. **b,** Core genome constructed with panaroo using previously published *E. coli* genomes from our laboratory (NCBI BioProject PRJNA819016). SNVs were extracted from the core genome alignment with snp-dists.

**Extended Data Figure 2.**
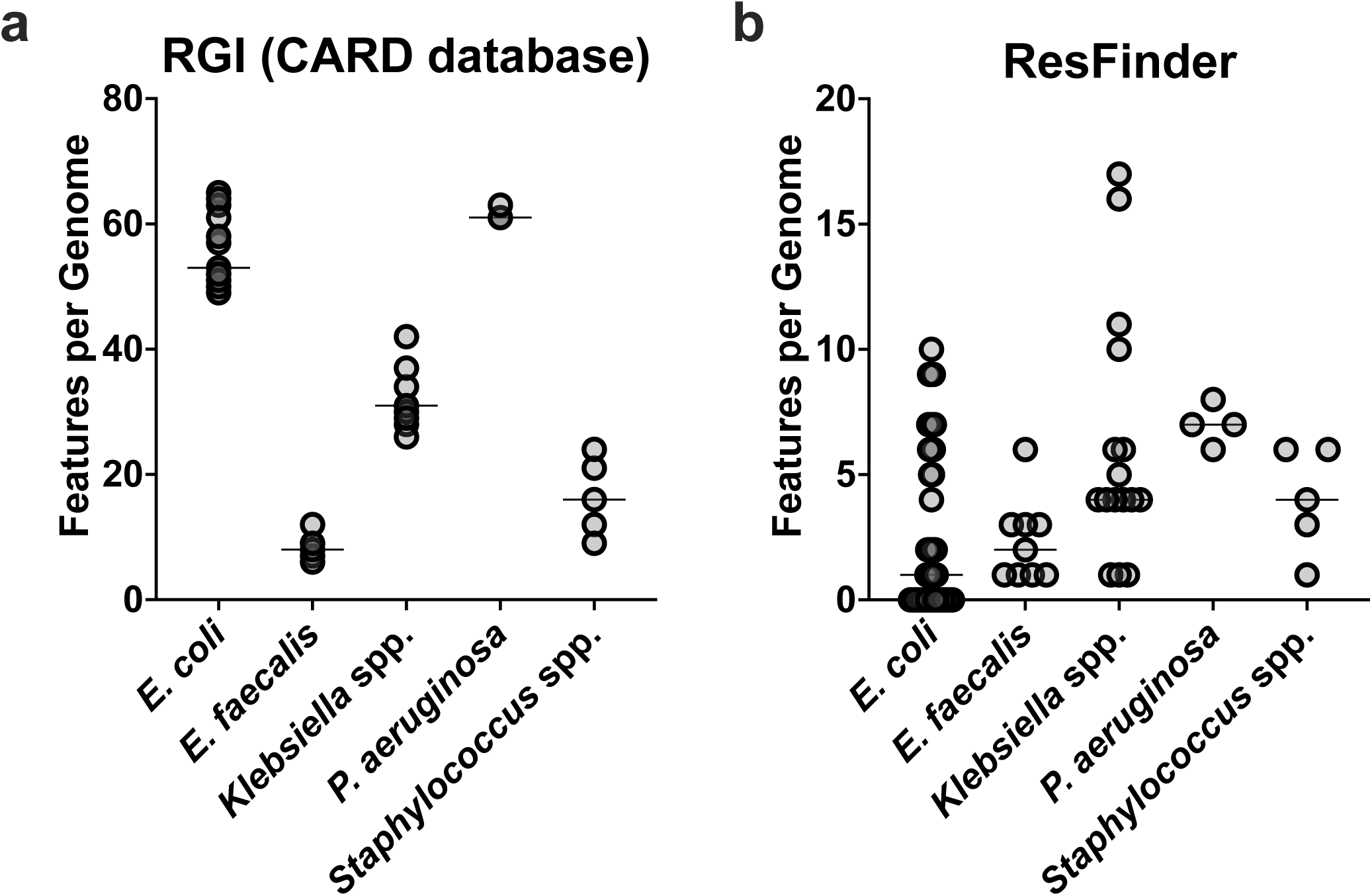
Frequency of antimicrobial resistance genes in bacterial genome assemblies by taxa. **a,** Distribution of antimicrobial resistance genes (ARGs) and stress response genes detected across bacterial taxa using the Resistance Gene Identifier (RGI) tool based on the Comprehensive Antibiotic Resistance Database (CARD) database (version 4.0.0). **b,** Distribution of ARGs detected by ResFinder (version 4.6.0). For both panels, the horizontal line depicts the median. Per genome results are detailed in Supplementary Table 5.

**Extended Data Figure 3.**
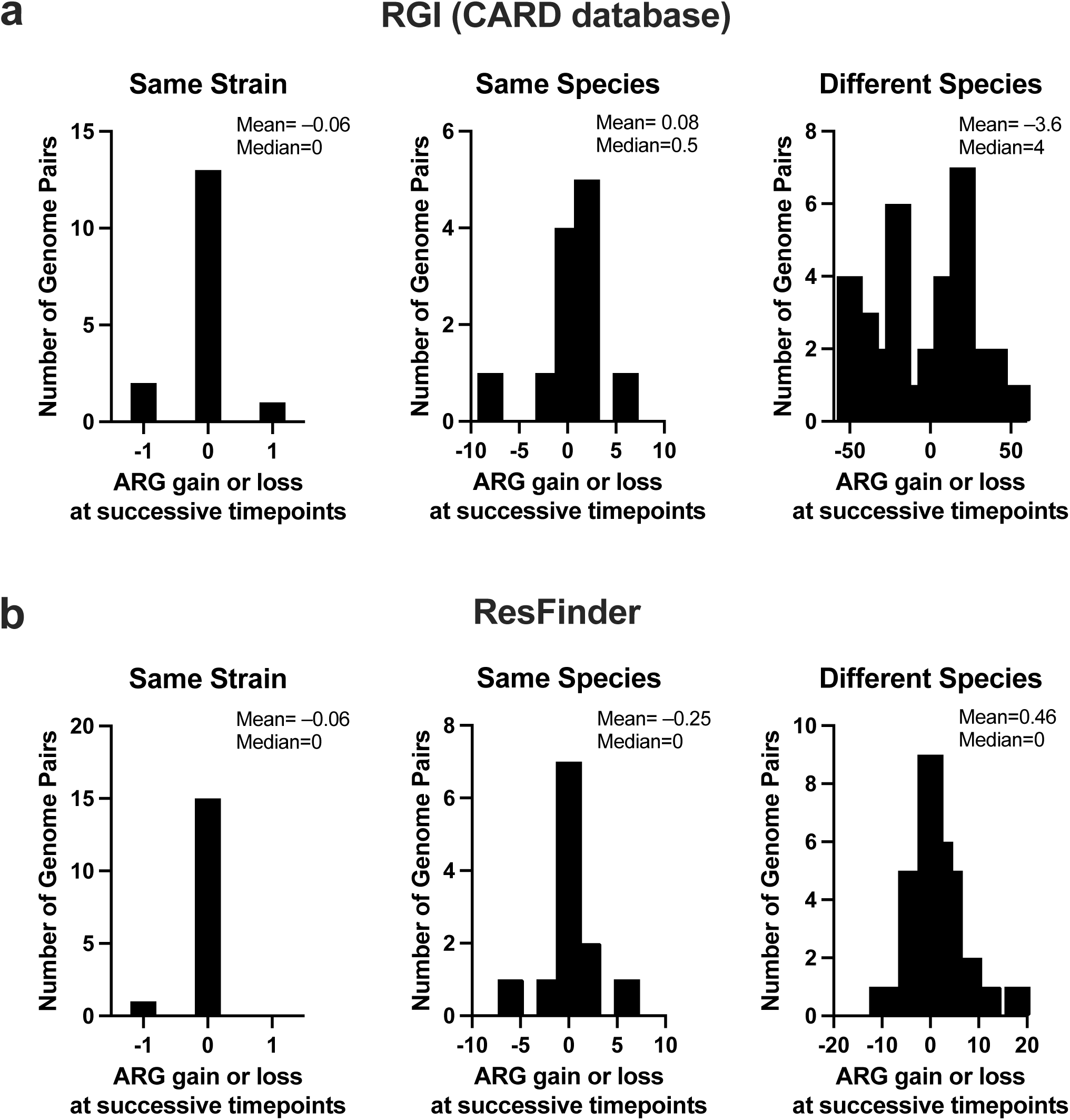
Comparison of antimicrobial resistance genes between successive samples from the same patients. Change in antimicrobial resistance gene (ARG) and stress response gene content between successive bacterial genomes from the same patients. ARGs and stress response genes identified by the Resistance Gene Identifier (RGI) tool based on the Comprehensive Antibiotic Resistance Database (CARD) database (version 4.0.0). **b,** Change in ARGs identified by ResFinder between successive bacterial genomes from the same patients. For both panels **a** and **b**, the horizontal axis depicts the differential in gene content between successive samples, and the vertical axis represents the number of genome pairs within the frequency bin.

**Extended Data Figure 4.**
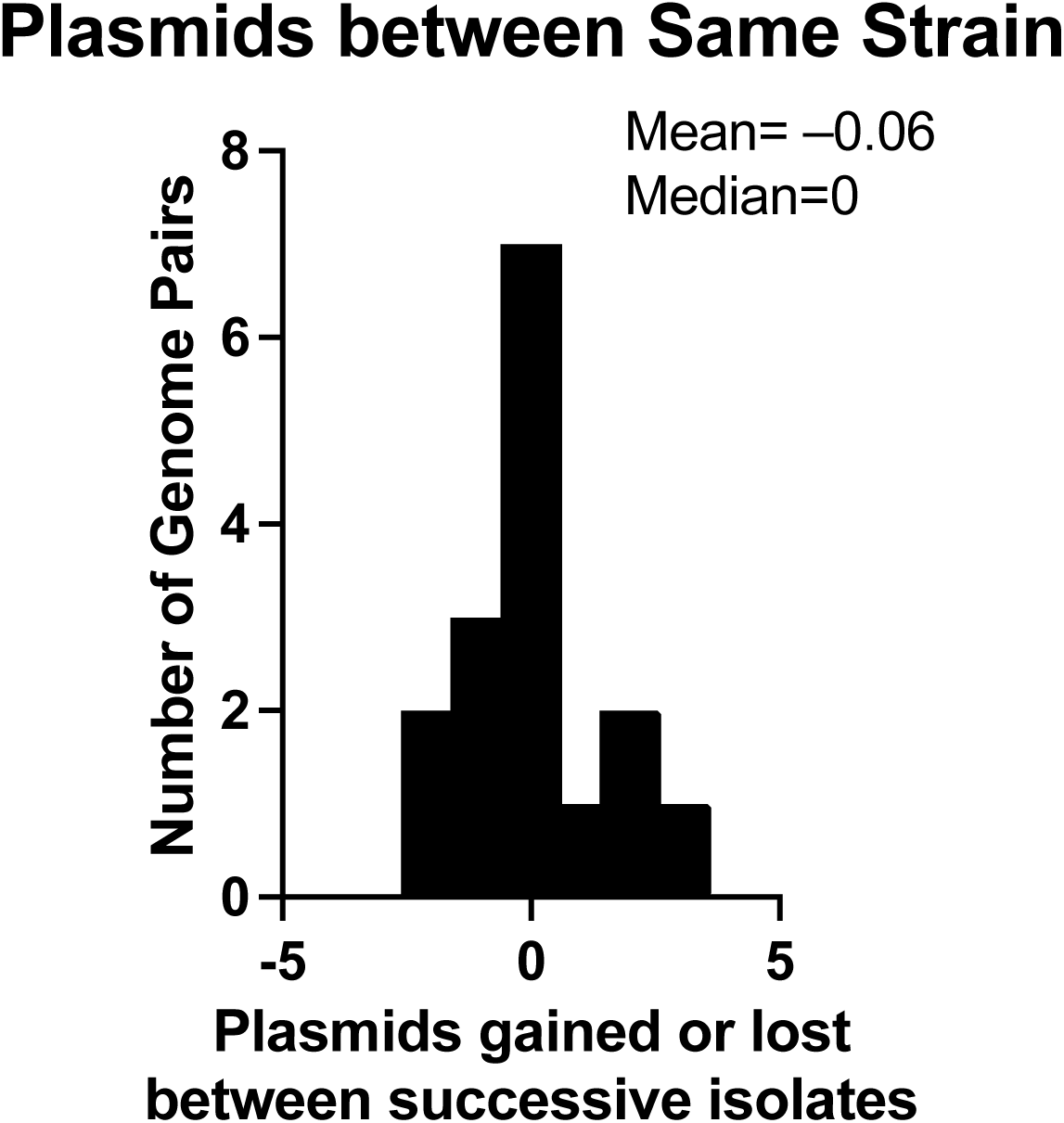
Plasmid content between successive samples of the same strain. Change in plasmid content between successive bacterial genomes from the same strain within individual patients. Plasmids were predicted from draft genome assemblies with MOB-suite.

**Extended Data Figure 5.**
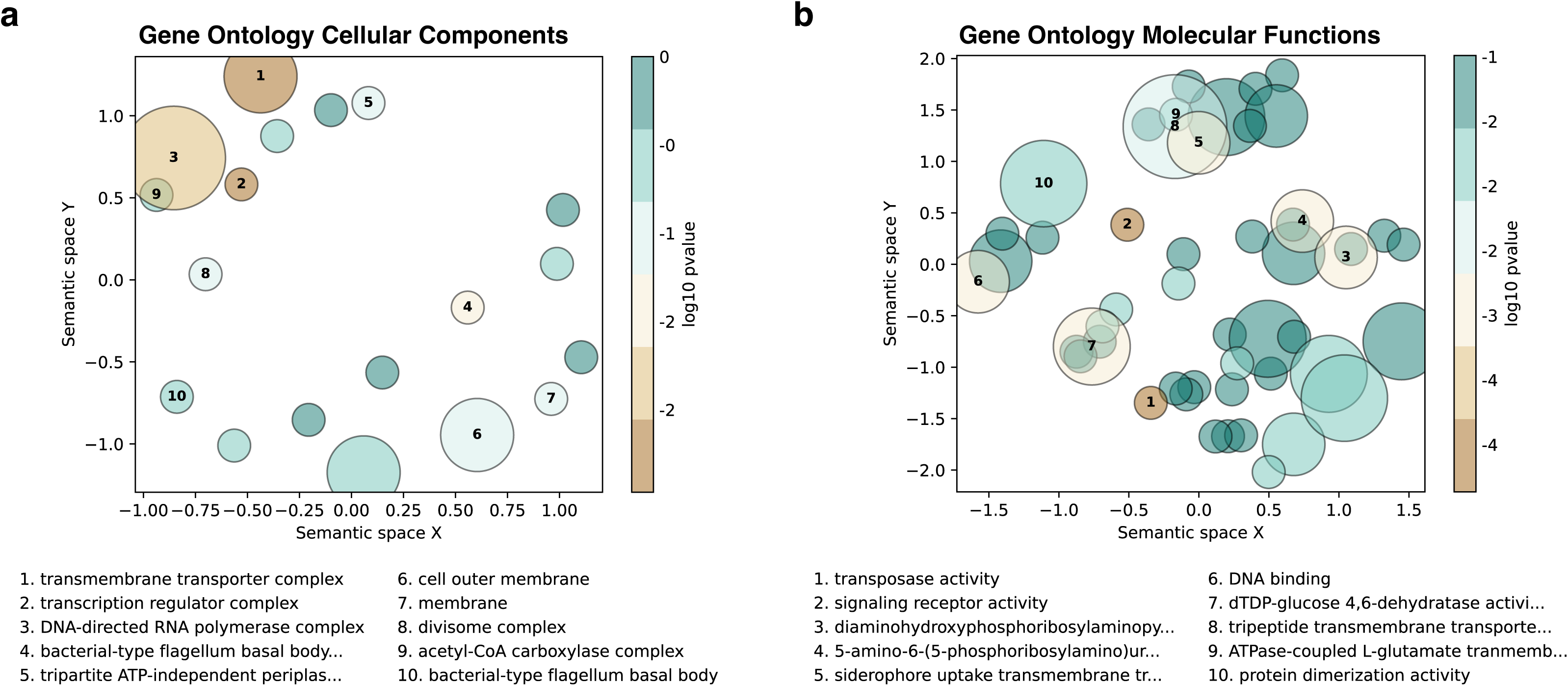
Gene ontology clustering from mutations. Cellular component (**a**) and molecular function (**b**) GO terms were clustered by semantic similarity with GO-Figure! ^104^. The size of the bubble is proportional to the number of genes associated with the particular semantic cluster. The color represents the Fisher exact test P-value from GO overrepresentation analysis (**Supplementary Table 9**).

**Extended Data Figure 6.**
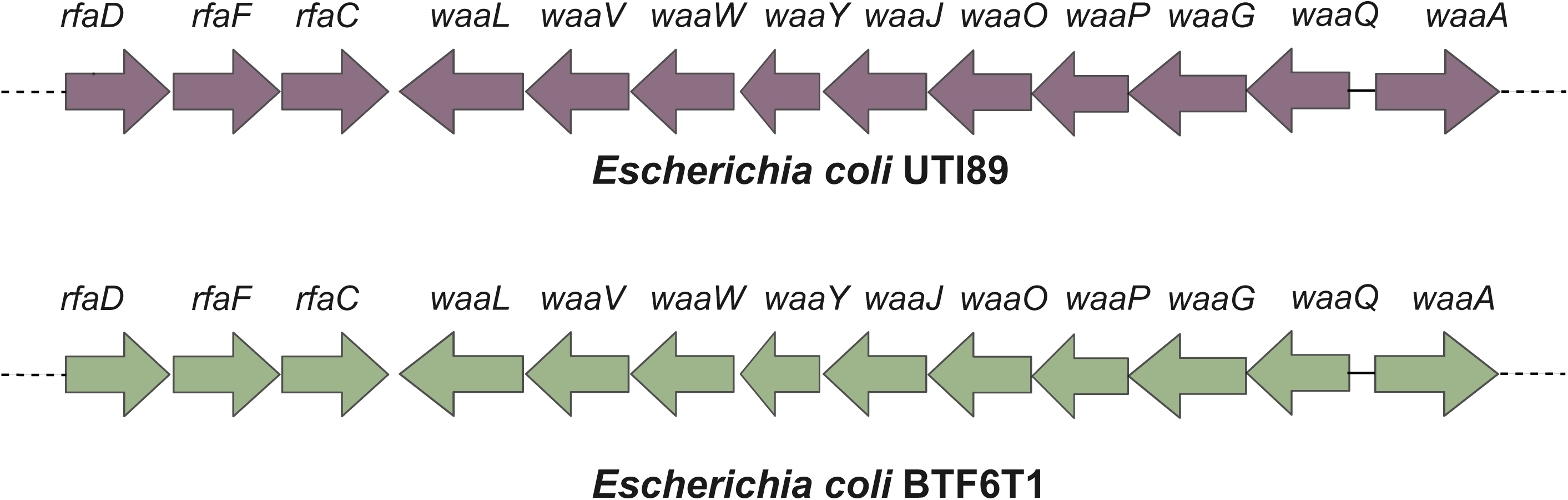
Operon organization of *waa* (formerly, *rfa*) locus in UTI89 and BTF6T1.

**Extended Data Figure 7.**
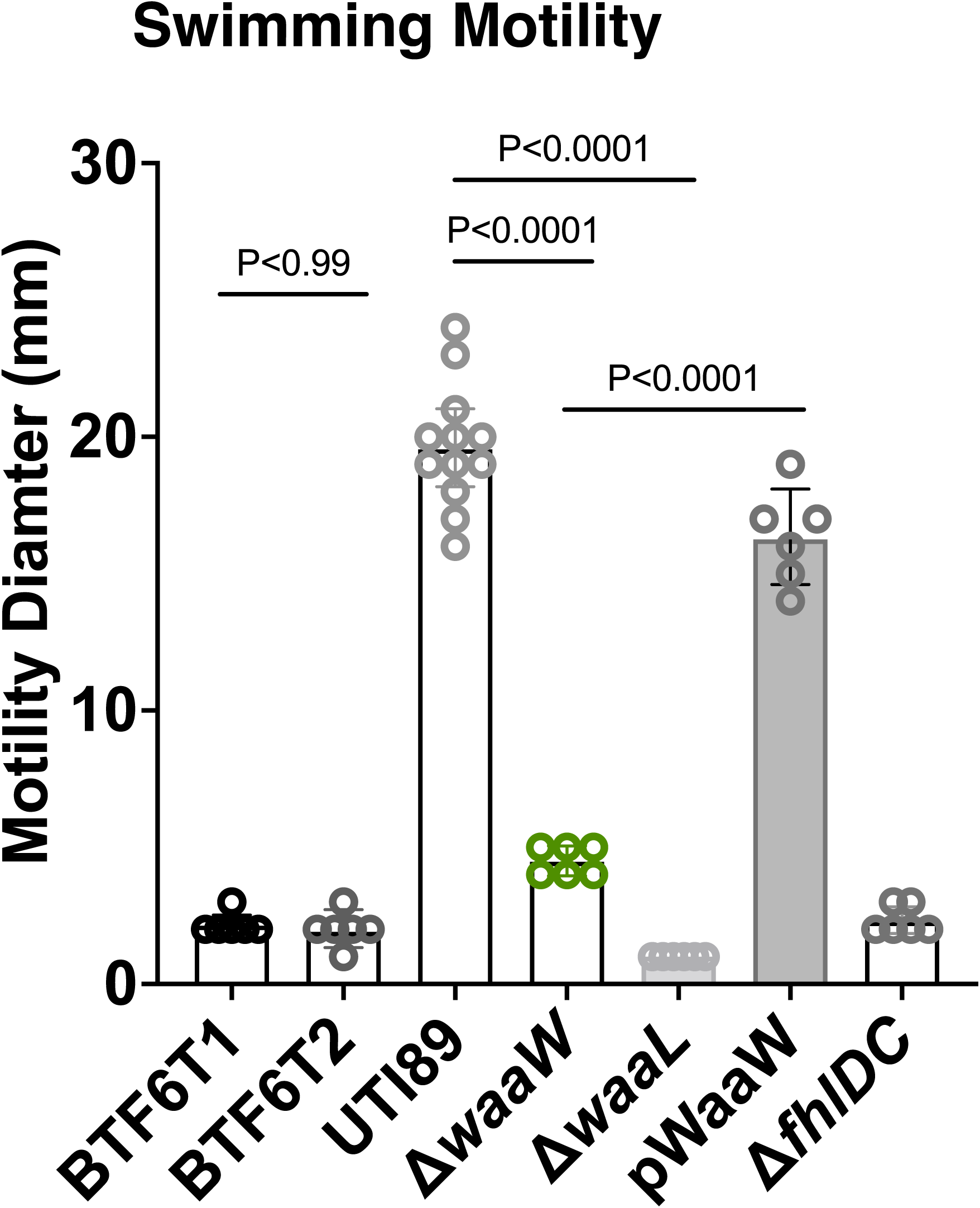
Soft agar swimming motility of clinical isolates BTF6T1, BTF6T2, UTI89, and its isogenic mutants *fhlDC, waaW,* and *waaL*. Complementation of the *waaW* deletion was conducted with the plasmid pTRC99a to produce pWaaW. UTI89Δ*fhlDC* is a negative transcriptional regulator of flagella and serves as a negative control. The bar height is the geometric mean, and the error bars depict the 95% CI. P values were calculated with non-parametric Kruskal–Wallis test with two-sided Dunn’s *post hoc* test for multiple comparisons.

**Extended Data Figure 8.**
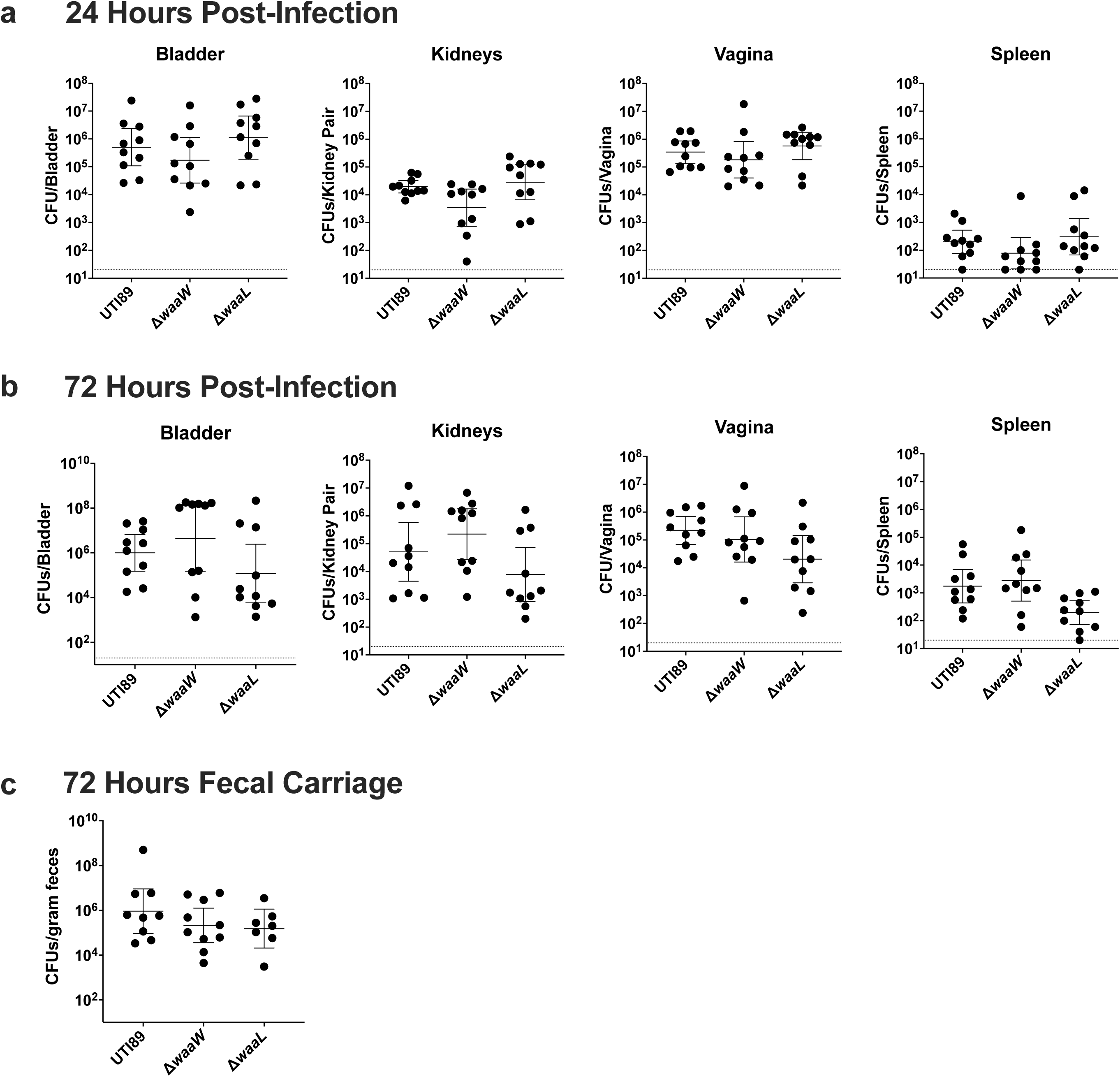
Loss of O-antigen does not impair bacterial fitness during an acute murine model of UTI. **a,** Bacterial burden in organs harvested 24-hours after transurethral inoculation with UTI89, UTI89Δ*waaW,* or UTI89Δ*waaL*. **b,** Bacterial burden in organs harvested 72-hours after transurethral inoculation with UTI89, UTI89Δ*waaW,* or UTI89Δ*waaL*. For panels **a** and **b**, the horizontal dashed line represents the limit of detection (20 CFUs per organ). **c,** Fecal bacterial counts of UTI89, UTI89Δ*waaW,* or UTI89Δ*waaL* 72-hours after transurethral inoculation. For all panels, the horizontal line depicts the geometric mean, and the error bars depict the 95% confidence interval. Statistical analysis was conducted with non-parametric Kruskal–Wallis test with two-sided Dunn’s *post hoc* test for multiple comparisons. No significant statistical differences were detected.

**Extended Data Figure 9.**
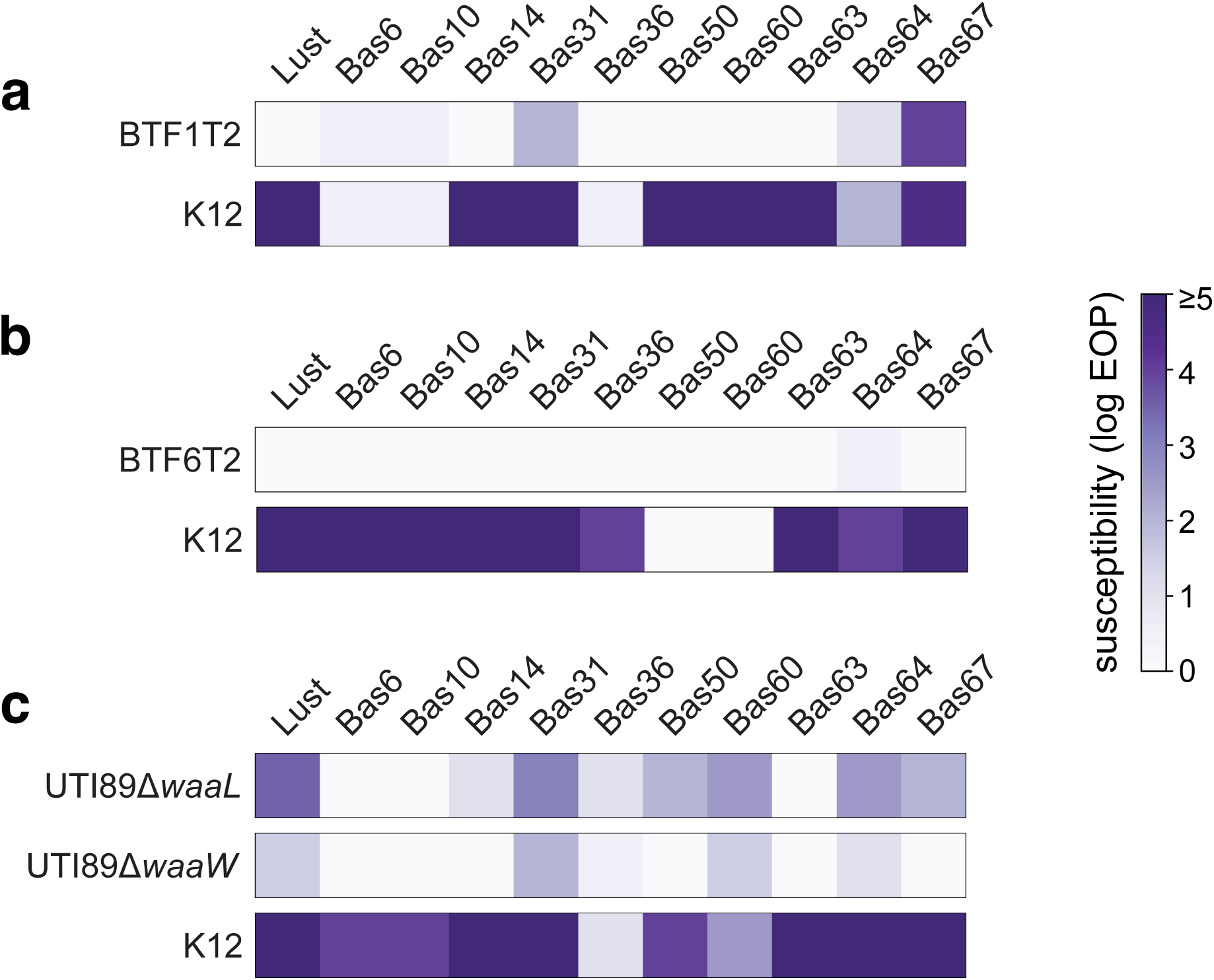
Phage susceptibility of UTI89 and its O-antigen biosynthesis mutants and select longitudinal *E. coli* isolates. Efficiency of plaquing (EOP) data against a panel of phages for the indicated bacterial strains compared to the corresponding strain BTF1T1 (**a**), BTF6T1 (**b**), and UTI89 (**c**). EOP values are displayed according to the legend on the right.

**Supplementary Table 1. Cohort characteristics.**

**Supplementary Table 2. Specimen metadata for 16S rRNA sequencing.**

**Supplementary Table 3. Bacterial genome accession and assembly metrics.**

**Supplementary Table 4. Antimicrobial susceptibility of sequenced colonies.**

**Supplementary Table 5. Antimicrobial resistance gene content of genome assemblies.**

**Supplementary Table 6. Predicted plasmids from genome assemblies.**

**Supplementary Table 7. Strains shared within individual patients.**

**Supplementary Table 8. Mutations from strains shared within individual patients.**

**Supplementary Table 9. Gene ontology overrepresentation analysis.**

**Supplementary Table 10. Bacterial strains and plasmids used for cloning.**

**Supplementary Table 11. Primers for cloning *waa* mutants.**

## Notes

### Competing Interest Statement

The authors have declared no competing interest.

## REFERENCES

1. Manack, A. et al. Epidemiology and healthcare utilization of neurogenic bladder patients in a US claims database. Neurourol Urodyn 30, (2011).

2. Kotkin, L. & Milam, D. F. Evaluation and management of the urologic consequences of neurologic disease. Tech Urol 2, (1996).

3. Ginsberg, D. The epidemiology and pathophysiology of neurogenic bladder. American Journal of Managed Care 19, (2013).

4. Mitchell, L. E. et al. Spina bifida. Lancet 364, (2004).

5. Copp, A. J., et al. Spina bifida. Nat Rev Dis Primers 1, (2015).

6. Snow-Lisy, D. C., Yerkes, E. B. & Cheng, E. Y. Update on Urological Management of Spina Bifida from Prenatal Diagnosis to Adulthood. Journal of Urology 194, 288–296 (2015).

7. Gebretsadik, T. et al. Rates of hospitalization for urinary tract infections among medicaid-insured individuals by spina bifida status, Tennessee 2005–2013. Disabil Health J 13, (2020).

8. Haudebert, C. et al. Risk factors for upper urinary tract deterioration in adult patients with spina bifida. World J Urol 41, 1187–1192 (2023).

9. Peyronnet, B. et al. Urologic Disorders are Still the Leading Cause of In-hospital Death in Patients With Spina Bifida. Urology 137, 200–204 (2020).

10. Armour, B. S. et al. Hospitalization for urinary tract infections and the quality of preventive health care received by people with spina bifida. Disabil Health J 2, 145–152 (2009).

11. Forster, C. S. & Wang, J. Symptom- and urinalysis-based approach to diagnosing urinary tract infections in children with neuropathic bladders. Pediatric Nephrology 35, 807–814 (2020).

12. Forster, C. S., Kowalewski, N. N., Atienza, M., Reines, K. & Ross, S. Defining Urinary Tract Infections in Children With Spina Bifida: A Systematic Review. Hospital Pediatrics vol. 11 1280– 1286 Preprint at 10.1542/hpeds.2021-005934 (2021).

13. Gupta, K. et al. International clinical practice guidelines for the treatment of acute uncomplicated cystitis and pyelonephritis in women: A 2010 update by the Infectious Diseases Society of America and the European Society for Microbiology and Infectious Diseases. Clinical Infectious Diseases 52, (2011).

14. Thompson, L. R. et al. A communal catalogue reveals Earth’s multiscale microbial diversity. Nature 551, 457–463 (2017).

15. Callahan, B. J. et al. DADA2: High-resolution sample inference from Illumina amplicon data. Nat Methods 13, 581–583 (2016).

16. Quast, C. et al. The SILVA ribosomal RNA gene database project: Improved data processing and web-based tools. Nucleic Acids Res 41, (2013).

17. Davis, N. M., Proctor, Di. M., Holmes, S. P., Relman, D. A. & Callahan, B. J. Simple statistical identification and removal of contaminant sequences in marker-gene and metagenomics data. Microbiome 6, (2018).

18. Karstens, L. et al. Controlling for Contaminants in Low-Biomass 16S rRNA Gene Sequencing Experiments. mSystems 4, (2019).

19. Mallick, H. et al. Multivariable association discovery in population-scale meta-omics studies. PLoS Comput Biol 17, (2021).

20. Chan, J. Z. M., Halachev, M. R., Loman, N. J., Constantinidou, C. & Pallen, M. J. Defining bacterial species in the genomic era: Insights from the genus Acinetobacter. BMC Microbiol 12, (2012).

21. Rodriguez-R, L. M. et al. An ANI gap within bacterial species that advances the definitionsdefinitionsof intra-species units. mBio 15, (2024).

22. Viver, T. et al. Towards estimating the number of strains that make up a natural bacterial population. Nat Commun 15, (2024).

23. Yan, Y., Nguyen, L. H., Franzosa, E. A. & Huttenhower, C. Strain-level epidemiology of microbial communities and the human microbiome. Genome Medicine vol. 12 Preprint at 10.1186/s13073-020-00765-y (2020).

24. Thänert, R. et al. Persisting uropathogenic Escherichia coli lineages show signatures of niche-specific within-host adaptation mediated by mobile genetic elements. Cell Host Microbe 30, 1034–1047.e6 (2022).

25. Grote, A. et al. Persistent Salmonella infections in humans are associated with mutations in the BarA/SirA regulatory pathway. Cell Host Microbe 32, 79–92.e7 (2024).

26. Olm, M. R. et al. inStrain profiles population microdiversity from metagenomic data and sensitively detects shared microbial strains. Nat Biotechnol 39, 727–736 (2021).

27. van Dijk, L. R. et al. StrainGE: a toolkit to track and characterize low-abundance strains in complex microbial communities. Genome Biol 23, (2022).

28. Kim, D. D. et al. Contaminated drinking water facilitates Escherichia coli strain-sharing within households in urban informal settlements. Nat Microbiol (2025) doi:10.1038/s41564-025-01986-w.

29. Jain, C., Rodriguez-R, L. M., Phillippy, A. M., Konstantinidis, K. T. & Aluru, S. High throughput ANI analysis of 90K prokaryotic genomes reveals clear species boundaries. Nat Commun 9, (2018).

30. Eberly, A. R. et al. Defining a Molecular Signature for Uropathogenic versus Urocolonizing Escherichia coli: The Status of the Field and New Clinical Opportunities. J Mol Biol 432, 786–804 (2020).

31. Bermudez, T. A. et al. Raising the alarm: fosfomycin resistance associated with non-susceptible inner colonies imparts no fitness cost to the primary bacterial uropathogen. Antimicrob Agents Chemother (2023) doi:10.1128/aac.00803-23.

32. Morales, G. et al. The Role of Mobile Genetic Elements in Virulence Factor Carriage from Symptomatic and Asymptomatic Cases of Escherichia coli Bacteriuria. Microbiol Spectr 11, (2023).

33. Green, H. D., et al. Intra-strain colony biofilm heterogeneity in uropathogenic Escherichia coli and the effect of the NlpI lipoprotein. Biofilm 8, (2024).

34. Tonkin-Hill, G. et al. Producing polished prokaryotic pangenomes with the Panaroo pipeline. Genome Biol 21, (2020).

35. Torsten Seemann. snp-dists. https://github.com/tseemann/snp-distsdoi:10.5281/zenodo.1411986.

36. Ghalayini, M. et al. Evolution of a dominant natural isolate of Escherichia coli in the human gut over the course of a year suggests a neutral evolution with reduced effective population size. Appl Environ Microbiol 84, (2018).

37. Stoesser, N. et al. Evolutionary history of the global emergence of the Escherichia coli epidemic clone ST131. mBio 7, (2016).

38. Von Mentzer, A. et al. Identification of enterotoxigenic Escherichia coli (ETEC) clades with long-term global distribution. Nat Genet 46, (2014).

39. Weinroth, M. D. et al. Rates of evolutionary change of resident Escherichia coli O157:H7 differ within the same ecological niche. BMC Genomics 23, (2022).

40. Mills, E. G. et al. A one-year genomic investigation of Escherichia coli epidemiology and nosocomial spread at a large US healthcare network. Genome Med 14, (2022).

41. Ludden, C. et al. Defining nosocomial transmission of Escherichia coli and antimicrobial resistance genes: a genomic surveillance study. Lancet Microbe 2, (2021).

42. Jia, B. et al. CARD 2017: Expansion and model-centric curation of the comprehensive antibiotic resistance database. Nucleic Acids Res 45, D566–D573 (2017).

43. Florensa, A. F., Kaas, R. S., Clausen, P. T. L. C., Aytan-Aktug, D. & Aarestrup, F. M. ResFinder – an open online resource for identification of antimicrobial resistance genes in next-generation sequencing data and prediction of phenotypes from genotypes. Microb Genom 8, (2022).

44. Bortolaia, V. et al. ResFinder 4.0 for predictions of phenotypes from genotypes. Journal of Antimicrobial Chemotherapy 75, (2020).

45. Zankari, E. et al. PointFinder: A novel web tool for WGS-based detection of antimicrobial resistance associated with chromosomal point mutations in bacterial pathogens. Journal of Antimicrobial Chemotherapy 72, (2017).

46. Robertson, J., Bessonov, K., Schonfeld, J. & Nash, J. H. E. Universal whole-sequence-based plasmid typing and its utility to prediction of host range and epidemiological surveillance. Microb Genom 6, (2020).

47. Robertson, J. & Nash, J. H. E. MOB-suite: software tools for clustering, reconstruction and typing of plasmids from draft assemblies. Microb Genom 4, (2018).

48. Barrick, J. E. et al. Identifying structural variation in haploid microbial genomes from short-read resequencing data using breseq. BMC Genomics 15, (2014).

49. Deatherage, D. E. & Barrick, J. E. Identification of mutations in laboratory-evolved microbes from next-generation sequencing data using breseq. Methods in Molecular Biology 1151, (2014).

50. Murphy, K. C. & Campellone, K. G. Lambda Red-mediated recombinogenic engineering of enterohemorrhagic and enteropathogenic E. coli. BMC Mol Biol 4, 1–12 (2003).

51. Cherepanov, P. P. & Wackernagel, W. Gene disruption in Escherichia coli: Tc R and Km R cassettes with the option of Flp-catalyzed excision of the antibiotic-resistance determinant (Antibiotic-resistance markers; endA; Flp recombinase; FRT site; gene replacement mutant; site-specific recombination; transposon). Gene 158, 9–14 (1995).

52. Datsenko, K. A. & Wanner, B. L. One-step inactivation of chromosomal genes in Escherichia coli K-12 using PCR products. Proceedings of the National Academy of Sciences 97, 6640–6645 (2000).

53. Mulvey, M. A., Schilling, J. D. & Hultgren, S. J. Establishment of a persistent Escherichia coli reservoir during the acute phase of a bladder infection. Infect Immun 69, 4572–4579 (2001).

54. Hadjifrangiskou, M. et al. Transposon mutagenesis identifies uropathogenic Escherichia coli biofilm factors. J Bacteriol 194, 6195–6205 (2012).

55. Davis, M. R. & Goldberg, J. B. Purification and visualization of lipopolysaccharide from gram-negative bacteria by hot aqueous-phenol extraction. Journal of Visualized Experiments (2012) doi:10.3791/3916.

56. Kulikov, E. E., Golomidova, A. K., Prokhorov, N. S., Ivanov, P. A. & Letarov, A. V. High-throughput LPS profiling as a tool for revealing of bacteriophage infection strategies. Sci Rep 9, (2019).

57. Fomsgaard, A., Freudenberg, M. A. & Galanos, C. Modification of the silver staining technique to detect lipopolysaccharide in polyacrylamide gels. J Clin Microbiol 28, (1990).

58. Martinez, J. J., Mulvey, M. A., Schilling, J. D., Pinkner, J. S. & Hultgren, S. J. Type 1 pilus-mediated bacterial invasion of bladder epithelial cells. EMBO Journal 19, 2803–2812 (2000).

59. Brannon, J. R. et al. Invasion of vaginal epithelial cells by uropathogenic Escherichia coli. Nat Commun (2020) doi:10.1038/s41467-020-16627-5.

60. Chaturvedi, K. S. et al. Cupric yersiniabactin is a virulence-associated superoxide dismutase mimic. ACS Chem Biol 9, 551–561 (2014).

61. Saenkham, P., Ritter, M., Donati, G. L. & Subashchandrabose, S. Copper primes adaptation of uropathogenic Escherichia coli to superoxide stress by activating superoxide dismutases. PLoS Pathog 16, 1–22 (2020).

62. Hung, C. S., Dodson, K. W. & Hultgren, S. J. A murine model of urinary tract infection. Nat Protoc 4, 1230–1243 (2009).

63. Hannan, T. J., Mysorekar, I. U., Hung, C. S., Isaacson-Schmid, M. L. & Hultgren, S. J. Early severe inflammatory responses to uropathogenic E. coli predispose to chronic and recurrent urinary tract infection. PLoS Pathog 6, 29–30 (2010).

64. Forde, B. M. et al. Population dynamics of an Escherichia coli ST131 lineage during recurrent urinary tract infection. Nat Commun 10, 1–10 (2019).

65. Thänert, R. et al. Comparative genomics of antibiotic-resistant uropathogens implicates three routes for recurrence of urinary tract infections. mBio 10, 1–16 (2019).

66. Luo, Y. et al. Similarity and divergence of phylogenies, antimicrobial susceptibilities, and virulence factor profiles of Escherichia coli isolates causing recurrent urinary tract infections that persist or result from reinfection. J Clin Microbiol 50, (2012).

67. Russo, T. A., Stapleton, A., Wenderoth, S., Hooton, T. M. & Stamm, W. E. Chromosomal restriction fragment length polymorphism analysis of escherichia coli strains causing recurrent urinary tract infections in young women. Journal of Infectious Diseases 172, (1995).

68. Skjøt-Rasmussen, L., et al. Persisting clones of Escherichia coli isolates from recurrent urinary tract infection in men and women. Journal of Medical Microbiology vol. 60 Preprint at 10.1099/jmm.0.026963-0 (2011).

69. Brannon, J. R. et al. Mapping niche-specific two-component system requirements in uropathogenic Escherichia coli. Microbiol Spectr 12, (2024).

70. Patzer, S. I., Albrecht, R., Braun, V. & Zeth, K. Structural and mechanistic studies of pesticin, a bacterial homolog of phage lysozymes. Journal of Biological Chemistry 287, (2012).

71. Whitfield, C. et al. Assembly of the R1-type core oligosaccharide of Escherichia coli lipopolysaccharide. J Endotoxin Res 5, (1999).

72. Amor, K. et al. Distribution of core oligosaccharide types in lipopolysaccharides from Escherichia coli. Infect Immun 68, (2000).

73. Raetz, C. R. H. & Whitfield, C. Lipopolysaccharide endotoxins. Annual Review of Biochemistry vol. 71 Preprint at 10.1146/annurev.biochem.71.110601.135414 (2002).

74. Leipold, M. D., Vinogradov, E. & Whitfield, C. Glycosyltransferases involved in biosynthesis of the outer core region of Escherichia coli lipopolysaccharides exhibit broader substrate specificities than is predicted from lipopolysaccharide structures. Journal of Biological Chemistry 282, (2007).

75. Romeyer Dherbey, J., Parab, L., Gallie, J. & Bertels, F. Stepwise Evolution of E. coli C and φX174 Reveals Unexpected Lipopolysaccharide (LPS) Diversity. Mol Biol Evol 40, (2023).

76. Heinrichs, D. E., Yethon, J. A., Amor, P. A. & Whitfield, C. The assembly system for the outer core portion of R1- and R4-type lipopolysaccharides of Escherichia coli: The R1 core-specific β-glucosyltransferase provides a novel attachment site for O-polysaccharides. Journal of Biological Chemistry 273, (1998).

77. Laekas-Hameder, M. & Daigle, F. Only time will tell: lipopolysaccharide glycoform and biofilm-formation kinetics in Salmonella species and Escherichia coli. J Bacteriol 206, e0031824 (2024).

78. Inoue, T. et al. Genome-wide screening of genes required for swarming motility in Escherichia coli K-12. J Bacteriol 189, 950–957 (2007).

79. Qin, J., Hong, Y., Morona, R. & Totsika, M. O antigen biogenesis sensitises Escherichia coli K-12 to bile salts, providing a plausible explanation for its evolutionary loss. PLoS Genet 19, (2023).

80. Browning, D. F., Hobman, J. L. & Busby, S. J. W. Laboratory strains of Escherichia coli K-12: things are seldom what they seem. Microb Genom 9, (2023).

81. Stevenson, G. et al. Structure of the O antigen of Escherichia coli K-12 and the sequence of its rfb gene cluster. J Bacteriol 176, (1994).

82. Liu, D. & Reeves, P. R. Escherichia coli K12 regains its O antigen. Microbiology (N Y) 140, (1994).

83. Fivenson, E. M. et al. A role for the Gram-negative outer membrane in bacterial shape determination. Proc Natl Acad Sci U S A 120, (2023).

84. Browning, D. F. et al. Laboratory adapted Escherichia coli K-12 becomes a pathogen of Caenorhabditis elegans upon restoration of O antigen biosynthesis. Mol Microbiol 87, (2013).

85. Jorgenson, M. A., Kannan, S., Laubacher, M. E. & Young, K. D. Dead-end intermediates in the enterobacterial common antigen pathway induce morphological defects in Escherichia coli by competing for undecaprenyl phosphate. Mol Microbiol 100, (2016).

86. Schooley, R. T. et al. Development and use of personalized bacteriophage-based therapeutic cocktails to treat a patient with a disseminated resistant Acinetobacter baumannii infection. Antimicrob Agents Chemother 61, (2017).

87. Maffei, E. et al. Systematic exploration of Escherichia coli phage-host interactions with the BASEL phage collection. PLoS Biol 19, (2021).

88. Humolli, D. et al. Completing the BASEL phage collection to unlock hidden diversity for systematic exploration of phage-host interactions. PLoS Biol 23, (2025).

89. Nagy, G. et al. Loss of regulatory protein RfaH attenuates virulence of uropathogenic Escherichia coli. Infect Immun 70, (2002).

90. Sarkar, S., Ulett, G. C., Totsika, M., Phan, M. D. & Schembri, M. A. Role of capsule and O antigen in the virulence of uropathogenic Escherichia coli. PLoS One 9, (2014).

91. Billips, B. K., Yaggie, R. E., Cashy, J. P., Schaeffer, A. J. & Klumpp, D. J. A live-attenuated vaccine for the treatment of urinary tract infection by uropathogenic escherichia coli. Journal of Infectious Diseases 200, (2009).

92. Coggon, C. F. et al. A novel method of serum resistance by Escherichia coli that causes urosepsis. mBio 9, (2018).

93. Burns, A. M. & Hull, S. I. Comparison of loss of serum resistance by defined lipopolysaccharide mutants and an acapsular mutant of uropathogenic Escherichia coli O75:K5. Infect Immun 66, (1998).

94. Burns, S. M. & Hull, S. I. Loss of resistance to ingestion and phagocytic killing by O- and K-mutants of a uropathogenic Escherichia coli O75:K5 strain. Infect Immun 67, (1999).

95. Lieberman, T. D. et al. Genetic variation of a bacterial pathogen within individuals with cystic fibrosis provides a record of selective pressures. Nat Genet 46, 82–87 (2014).

96. Lieberman, T. D. et al. Parallel bacterial evolution within multiple patients identifies candidate pathogenicity genes. Nat Genet 43, 1275–1280 (2011).

97. Poret, A., et al. De novo mutations mediate phenotypic switching in an opportunistic human lung pathogen. bioRxiv (2024) doi:10.1101/2024.02.06.579193.

98. Silva, I. N., et al. Long-Term Evolution of Burkholderia multivorans during a Chronic Cystic Fibrosis Infection Reveals Shifting Forces of Selection. mSystems 1, (2016).

99. Hassan, A. A., Coutinho, C. P. & Sá-Correia, I. Burkholderia cepacia Complex Species Differ in the Frequency of Variation of the Lipopolysaccharide O-Antigen Expression During Cystic Fibrosis Chronic Respiratory Infection. Front Cell Infect Microbiol 9, (2019).

100. Golubchik, T. et al. Within-Host Evolution of Staphylococcus aureus during Asymptomatic Carriage. PLoS One 8, (2013).

101. Marvig, R. L., Sommer, L. M., Molin, S. & Johansen, H. K. Convergent evolution and adaptation of Pseudomonas aeruginosa within patients with cystic fibrosis. Nat Genet 47, 57–64 (2015).

102. Raghuram, V. et al. Comparison of genomic diversity between single and pooled Staphylococcus aureus colonies isolated from human colonization cultures. Microb Genom 9, (2023).

103. Yu, G., Smith, D. K., Zhu, H., Guan, Y. & Lam, T. T. Y. ggtree: an r package for visualization and annotation of phylogenetic trees with their covariates and other associated data. Methods Ecol Evol 8, (2017).

104. Reijnders, M. J. M. F. & Waterhouse, R. M. Summary Visualizations of Gene Ontology Terms With GO-Figure! Frontiers in Bioinformatics 1, (2021).

105. O’Toole, Á., Aziz, A. & Maloney, D. Publication-ready single nucleotide polymorphism visualization with snipit. Bioinformatics 40, (2024).

