## Supplementary material for "Unmasking Pathogen Traits for Chronic Colonization in Neurogenic Bladder Patients": Cohort characteristics.

| **Participants (n)** | **77** |
| --- | --- |
| **Age, years** **(mean, [range])** | 7.1 [0-27.4] |
| **Sex, n (%)**  Male  Female | 41 (53.2)  36 (46.8) |
| **Race, n (%)**  White  Black  Asian | 68 (88.3)  6 (7.8)  3 (3.9) |
| **Type of Delivery, n (%)**  Vaginal Birth  Cesarean Section  Unknown | 12 (15.6)  61 (79.2)  4 (5.2) |
| **Type of Lesion, n (%)**  Myelomeningocele  Lipomyelomeningocele  Other | 70 (90.9)  4 (5.2)  3 (3.9) |
| **Bladder Management, n (%)**  Clean Intermittent Catheterization (CIC) | 44 (57.1) |
| **Antibiotic Use in Past 30 days, n (%)** | 23 (29.9) |
| **Any History of UTI, n (%)** | 49 (63.6) |
