## Supplementary material for "Unmasking Pathogen Traits for Chronic Colonization in Neurogenic Bladder Patients": Bacterial strains and plasmids used for cloning.

| **Strain or Plasmid** | **Description** | **Source** |
| --- | --- | --- |
| UTI89 | *E. coli* cystitis isolate | PMID: 11402001 |
| UTI89Δ*waaW* | UDP-galactose:(galactosyl) LPS alpha 1,2-galactosyltransferase (UTI89_C4169) deletion mutant | This work |
| UTI89Δ*waaL* | O-antigen ligase (UTI89_C4167) deletion mutant | This work |
| UTI89Δ*flhDC* | Master regulator of flagellar biosynthesis (UTI89_C2094/C2095) deletion mutant | PMID: 21542868 |
| K12 MG1655 | *E. coli* laboratory reference strain (*wbbL* -) | PMID: 40409266 |
| pKD4 | Encodes FRT-flanked kanamycin resistance cassette | PMID: 10829079 |
| pKM208 | Encodes λ Red recombinase | PMID: 14672541 |
| pCP20 | Encodes temperature-sensitive FLP recombinase | PMID: 7789817 |
| pTRC99a | Empty vector | PMID: 3069586 |
| pWaaW | *waaW* complemented in pTRC99a under its native promoter | This work |
