## Supplementary material for "Unmasking Pathogen Traits for Chronic Colonization in Neurogenic Bladder Patients": Primers for cloning waa mutants.

| **Primer Name** | **Sequence (5′-3′)** | **Purpose** |
| --- | --- | --- |
| waaL_KO_F | ATGTCGTTTTGTTGGAATGAAATTAACTCTGGTGTCAAGTCTTTAATTCGTGTAGGCTGGAGCTGCTTC | *waaL* deletion |
| waaL_KO_R | CTAATAAACATTGGTCCGATTGTACTTTAAAATAAGCACAAAGCAATCATATGAATATCCTCCTTAG | *waaL* deletion |
| waaL_test_F | CTATATTCTCTGGAGGATGGTAAGAC | Test primer for *waaL* deletion |
| waaL_test_R | GGAATGAGTTGTCTCAATTGACAGC | Test primer for *waaL* deletion |
| waaW_KO_F | GCCCAAATAGTTTTCTTTTATACTTATTTAATTTGAATTTAATGAAGGTGTAGGCTGGAGCTGCTTC | *waaW* deletion |
| waaW_KO_R | CAACATGGATTTATTAGCTGAGAGTATTACTGAAGTCGCTGTCTCTGGGGCATATGAATATCCTCCTTAG | *waaW* deletion |
| waaW_test_F | GTACTCATCCTTAATTATTATTG | Test primer for *waaW* deletion |
| waaW_test_R | GGCAAAGCGTAAACCACAC | Test primer for *waaW* deletion |
| waaW_gibson_F | ACACAGGAAACAGACCATGGCATCCATGATTTTTATTATTGCCCAAATAG | Amplification of *waaW* from genomic DNA |
| waaW_gibson_R | TCTAGAGGATCCCCGGGTACCGTCACCTGGGTATTGCC | Amplification of *waaW* from genomic DNA |
| pTRC99a_mcs_F | TAATTCGTGTCGCTCAAGGC | Test primer for complementation with pTRC99a |
| pTRC99a_mcs_R | CCGCCAGGCAAATTCTGTTT | Test primer for complementation with pTRC99a |
