## Supplementary material for "Unmasking Pathogen Traits for Chronic Colonization in Neurogenic Bladder Patients": Strains shared within individual patients.

| **Reference Genome** | **Query Genome** | **Species** | **Days Between** |
| --- | --- | --- | --- |
| BTF1T1 | BTF1T2 | *Escherichia coli* | 437 |
| BTF2T1Ec | BTF2T2b | *Escherichia coli* | 399 |
| BTF6T1 | BTF6T2 | *Escherichia coli* | 79 |
| BTF15T-1 | BTF15T1 | *Klebsiella pneumoniae* | 811 |
| BTF24T1 | BTF24T2 | *Escherichia coli* | 411 |
| BTF28T-2 | BTF28T-1 | *Klebsiella pneumoniae* | 46 |
| BTF37T1 | BTF37T2 | *Escherichia coli* | 392 |
| BTF37T1b | BTF37T2b | *Enterococcus faecalis* | 392 |
| BTF46T1 | BTF46T2b | *Proteus mirabilis* | 364 |
| BTF60T-3 | BTF60T-1 | *Escherichia coli* | 305 |
| BTF60T-3 | BTF60T-2 | *Escherichia coli* | 134 |
| BTF60T1 | BTF60T2 | *Escherichia coli* | 343 |
| BTF69T-2 | BTF69T-1 | *Escherichia coli* | 178 |
| BTF71T-3 | BTF71T-2 | *Escherichia coli* | 126 |
| BTF74T-7 | BTF74T-6 | *Escherichia coli* | 69 |
| BTF75T-1 | BTF75T1 | *Escherichia coli* | 602 |
